# Double-edged swords: Anthracyclines inhibit -1 programmed ribosomal frameshifting and restrict HCoV-OC43 infection but show cytotoxicity

**DOI:** 10.64898/2026.03.08.709729

**Authors:** Daniel Scheller, Koushikul Islam, Karolis Vaitkevicius, Dmitriy Ignatov, Lisa Pettersson, Lena Lindgren, Niklas Arnberg, Jörgen Johansson

## Abstract

Human coronavirus OC43 (HCoV-OC43) constitutes one of the most common causes of the seasonal cold but can also cause severe disease among elderly and immuno-compromised. Currently, there are no approved antiviral drugs to combat HCoV-OC43 infection. Coronaviruses are positive-sense single-stranded RNA (+ssRNA) viruses and utilize -1 programmed ribosomal frameshifting (-1 PRF) to produce the correct stoichiometry of viral protein components. Due to its high conservation, the ribosomal frameshifting stimulation element (FSE) is a promising target for antiviral drug discovery. Aminoglycosides and anthracyclines interact with structured nucleic acids and some of these compounds affect -1 PRF in SARS-CoV-2. To get a comprehensive view of their putative efficacy, we examined all commercially available aminoglycosides and anthracyclines for their ability to affect -1 PRF in HCoV-OC43 and their potential to reduce HCoV-OC43 infection. Of 25 candidates tested, we identified 3 anthracyclines (Idarubicin, Nogalamycin and Daunorubicin) to reduce HCoV-OC43 -1 PRF using *in vitro* assays. We further demonstrate that the active anthracyclines, but not the inactive anthracycline, bind to the HCoV-OC43 FSE. The active anthracyclines did not significantly affect -1 PRF in SARS-CoV-2, suggesting differences in the interactions of the anthracyclines and FSEs, despite being relatively conserved in HCoV-OC43 and SARS-CoV-2 RNAs. Interestingly, these potent anthracyclines also significantly reduced HCoV-OC43 infection in human host cells. Although, the anthracyclines show some toxicity and introduce lesions in cellular DNA they could constitute an important scaffold for further antiviral development, with the aim to increase efficacy and reduce toxicity.

**Author summary:** The recent Covid-19 pandemic caused by SARS-CoV-2 showed the need for functional antivirals to immediately combat infections at the onset of future pandemics. In addition to SARS-CoV-2, four other human coronaviruses can cause mild to severe infection. Among those is HCoV-OC43, a coronavirus particularly troublesome for immunocompromised patients. Despite this clinical relevance, no approved antiviral therapies are currently available for the prevention or treatment of HCoV-OC43 infections.

In this work, we have identified a subset of anthracyclines (Idarubicin, Nogalamycin and Daunorubicin) that selectively and significantly reduce HCoV-OC43 infection. In contrast to a non-active anthracycline, the same anthracyclines that reduce infection also reduce ribosomal frameshifting and interact with a structured RNA in the viral RNA-genome. Interestingly, Idarubicin, a clinically used anticancer drug, shows selectivity for HCoV-OC43, but not for SARS-CoV-2 RNA. Our work demonstrates that anthracyclines could be important for pandemic preparedness and serve as attractive scaffolds when developing new antivirals targeting coronaviruses.

## 1. Introduction

Coronaviruses are single-stranded positive-sense RNA viruses, belonging to the order of *Nidovirales* and more specifically to the family *Coronaviridae* [1]. Seven coronaviruses (CoVs) infect humans [2]. Among them, human coronavirus OC43 (HCoV-OC43), HCoV-229E, HCoV-NL63 and HCoV-HKU1, also known as seasonal coronaviruses, cause upper-respiratory tract illnesses and are the cause of 15% - 30% of the seasonal common cold cases. Middle East respiratory syndrome coronavirus (MERS-CoV) and the severe acute respiratory syndrome coronaviruses (SARS-CoV, SARS-CoV-2) cause severe respiratory diseases, and together with other, non-human coronaviruses, these viruses are considered to be of high pandemic potential [3]. Covid-19 can also lead to long-covid, which is currently challenging to treat, cure and prevent [4]. The World Health Organization has declared eight public health emergencies of international concern, all caused by viruses and the majority of these resulting from zoonotic events [5]. Owing to the known plethora of viruses in nature [6] and the knowledge that climate change further increases the risk for zoonotic virus spillover from nature [7], the discovery and development of new antiviral compounds is critical for pandemic preparedness.

Small-molecule targeting of conserved RNA-structures is a strategy with great potential, especially for viruses harbouring a single-stranded RNA (ssRNA) genome. As a difference to linear RNA-sequences, RNA-structures are believed to be less prone of developing drug-resistance, since they have been evolutionarily retained by selective pressure [8]. For drug development, these structures should ideally be thermodynamically stable, fulfil a critical role in gene regulation and be specific to the pathogen [9]. Recently, the frameshifting stimulation element (FSE) dictating -1 programmed ribosomal frameshift (-1 PRF) was shown to be a promising target for antiviral drug discovery [10–16]. -1 PRF is a strategy used by many RNA viruses, such as CoVs to increase their coding capacity and also to regulate the ratio of different proteins [17–19]. It is a regulated translational recoding event in which the ribosome shifts by one nucleotide (-1) relative to the original reading frame, resulting in the synthesis of an alternative protein from the same mRNA template [20,21]. Mechanistically, -1 PRF relies on the FSE that comprises a slippery sequence followed downstream by a stable RNA structure, typically a pseudoknot, that together promote ribosomal pausing and induce a shift into the -1-reading frame [19,22]. The FSE is structurally highly conserved in the *Coronaviridae* family. In the subgroup of beta-coronavirus the FSE exhibits a three-stemmed pseudoknot structure preceded by a slippery site where the ribosome can slide back one base, thus changing the reading frame[23] (Figure 1).

**Figure 1.**
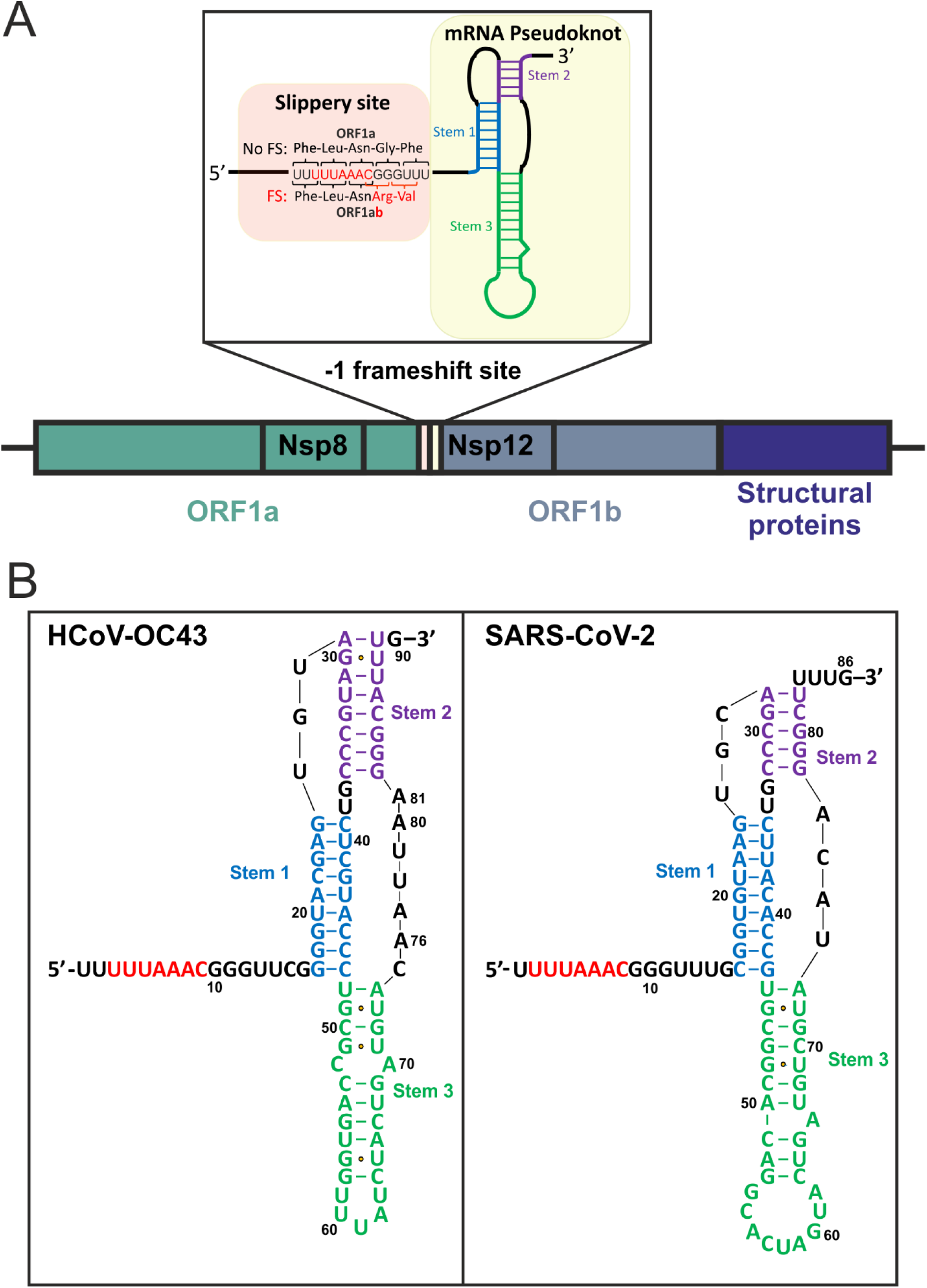
Genomic RNA organisation in coronaviruses. **(A)** Schematic representation of the genomic RNA organisation of coronaviruses with the -1 PRF site highlighted in the box. **(B)** Secondary RNA structure of the frameshift element of HCoV-OC43 and SARS-CoV-2.

Translation from the open reading frames (ORF) ORF1a and ORF1b generates most of the viral proteins. Expression of the RNA replication machinery of the virus, such as the RNA-dependent RNA polymerase and other RNA modifying enzymes, are encoded on ORF1b. Translation of the ORF1b requires -1 PRF (Figure 1). As a result, ORF1a and ORF1b are synthesized in a fine-tuned ratio with a 1.5 to 2-fold excess of ORF1a [24,25]. Consequently, disrupting the balance of ORF1a vs. ORF1b levels leads to severely crippled viral replication [12,15,26,27]. It could be hypothesized that molecules dysregulating -1 PRF could alter the balance thus affecting viral propagation. Aminoglycosides (Figure 2) are a class of naturally derived antibiotics that bind preferentially to structured RNA, including 16S ribosomal RNA in the bacterial decoding center, where they disrupt translational fidelity and induce miscoding [28]. Furthermore, they interact with other structured RNA elements such as ribozymes and RNase P [29]. The aminoglycoside geneticin has been shown to inhibit SARS-CoV-2 replication by interfering with -1 PRF, supporting the idea of targeting the FSE as an antiviral strategy [16]. Another interesting class of RNA-binding molecules are anthracyclines (Figure 2), known to bind non-coding RNAs [30], iron-responsive elements consisting of hairpin-loop structures, [31] and tRNAs [32]. Anthracyclines are clinically used mainly for cancer treatments [33,34], but have also been reported to inhibit viral replication in cells [35,36]. Members of this compound class suppress both infection of hepatitis B virus and of Ebola virus through activation of the innate immune response [37,38]. The anthracycline Aclarubicin (also known as Aclacinomycin) interferes with SARS-CoV-2 RNA-dependent RNA polymerase activity [35]. However, there is currently no evidence that anthracyclines inhibit -1 PRF by targeting structured RNA elements such as the FSE.

**Figure 2.**
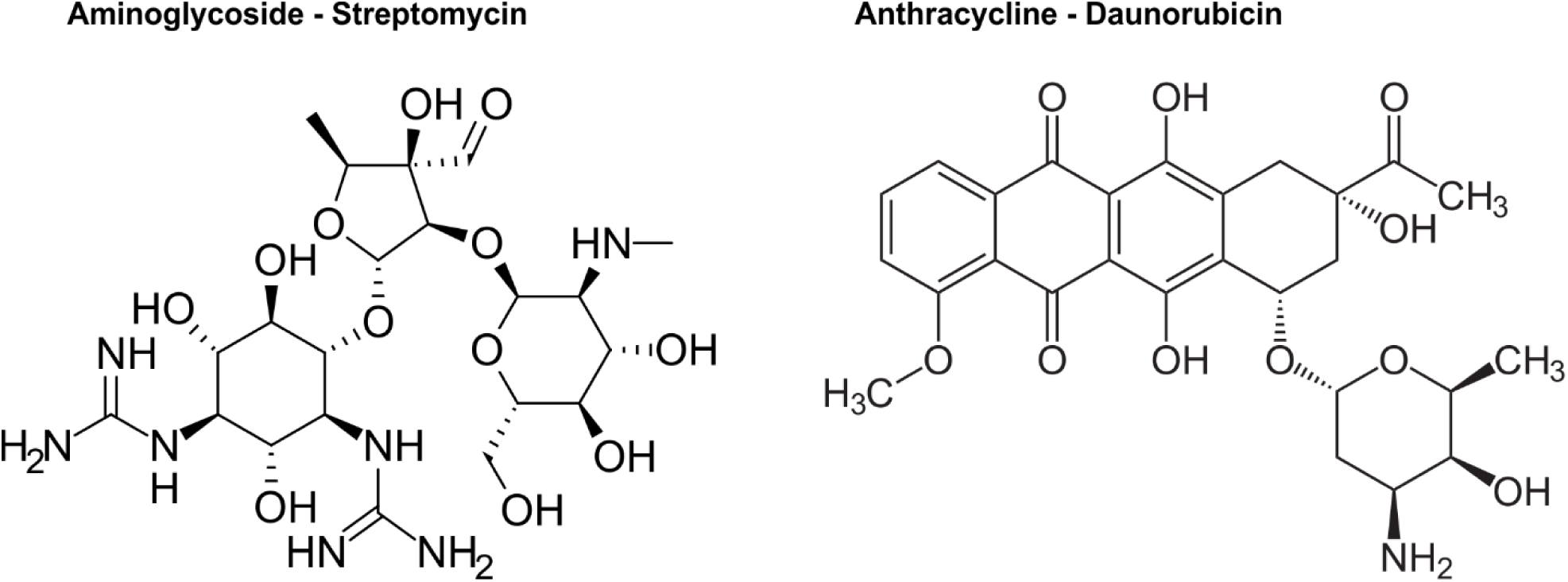
Chemical structure of the aminoglycoside Streptomycin and the anthracycline Daunorubicin as representative members of their respective compound class.

In this study, we investigated the potential of aminoglycosides and anthracyclines to influence -1 PRF in HCoV-OC43 and SARS-CoV-2. Using *in vitro* translation, fluorescence quenching and DMS-MaPseq, we show that a subset of anthracyclines can reduce -1 PRF by binding to the HCoV-OC43 FSE. These anthracyclines also substantially reduced HCoV-OC43 infectivity but also introduced DNA-lesions. Anthracyclines could thus prove important scaffolds for developing effective anti-HCoV-OC43 drugs, but efforts need to be taken to increase efficacy and decrease toxicity.

## 2. Material and Methods

### 2.1 Cells and viruses

#### Cell lines

HCT-8 cells were maintained in RPMI-1640 supplemented with 2 g NaHCO_3_/L, 10% fetal bovine serum (FBS, Cytiva) and 1% PEST (penicillin G (100 IU/mL) and streptomycin sulfate (100 μg/mL) combined) (Gibco) at 37°C and 5% CO_2_. Human angiotensin-converting enzyme 2 (ACE2)-expressing A549 (A549-ACE2) cells (provided by Professor Oscar Fernandez-Capetillo Laboratory, KI, SciLifeLab, Sweden), Calu-3 and VeroE6 cells were maintained in DMEM supplemented with 2 g NaHCO_3_/L, 10% FBS (VeroE6 5%) and 1%PeST at 37°C and 5% CO_2_.

#### Viruses

HCoV-OC43 (VR-1558, ATCC) were propagated in HCT-8 cells and titrated using focus-forming assay. The SARS-CoV-2 patient isolate SARS-CoV-2/01/human/2020/SWE; GenBank accession no. MT093571.1 was provided by the Public Health Agency of Sweden. The Delta variant (lineage B.1.617.2) strain SARS-CoV-2/Hu/DNK/SSI-H11/2021 (GenBank accession no. OM444216) and the Omicron BA.2 variant SARS-CoV-2/human/DNK/SSI-H53/2021(GenBank accession no. OM618106.1) were kindly provided by Dr. Charlotta Polacek Strandh, Virus & Microbiological Special Diagnostics, Statens Serum Institut, Copenhagen, Denmark). The SARS-CoV-2 XBB.1.9.2.5.1.3 variant SARS-CoV-2/hu/2023/SWE (GenBank accession no. PX643056) was collected from patient nasopharyngeal swab and isolated and propagated in Calu-3 cells in our laboratory.

### 2.2 Compounds

Nogalamycin, Pirarubicin, Mithramycin A and Aclacinomycin A have been purchased from Santa Cruz Biotechnology. Merafloxacin has been purchased from TargetMol Chemicals. All other compounds have been purchased from Sigma Aldrich. Compounds were dissolved in DMSO to a stock concentration of 10 mM and stored either at 4°C or -20°C according to manufacturers’ instruction.

### 2.3 Effect of compounds on HCoV-OC43 infection

Antiviral activity of 25 compounds against HCoV-OC43 virus was conducted using immunofluorescence-based assay as described previously [39]. Briefly, one day before infection, HCT-8 cells were seeded in 96 well plates (approximately 1.8 × 10^4^ cells/well) supplemented with cell maintaining media. Compounds were diluted to 1 μM and 5 μM final concentrations and mixed with approximately 1800 FFUs of HCoV-OC43 in RPMI containing 2% FBS, with the aim to have a multiplicity of infection (MOI) of 0.1. The cells were then infected with the virus and compound mixture and incubated for 16 h at 33°C with 5% CO_2_. HCoV-OC43 infection was assessed following incubation with primary rabbit polyclonal antibody directed against virus nucleoprotein (1:1000 dilution, 40643-T62, SinoBiological) and secondary anti-rabbit Alexa Fluor 647 antibody 1:1000, (A-31573, Invitrogen); cellular nuclei were stained with 0.1% DAPI for 10 min. Antibody-bound infection foci and cell nucleus were visualized and counted using cell imaging Cytation 5 microscope (Agilent BioTek).

### 2.4 IC_50_ value determination of anthracyclines compounds

The IC_50_ value of compounds was determined by immunofluorescence-based assay as described in the previous section. The individual infectious cell foci were quantified in response to compound treatment in a dose-dependent manner. Compounds were serially diluted in two-fold steps from 40 to 0.156 μM and mixed with HCoV-OC43 virus (MOI = 0.1) in RPMI-1640 containing 2% FBS and added to the HCT-8 cells. The plates were incubated for 16 h at 33°C in 5% CO_2_. Virus infection and number of cells in each well were quantified following antibody and DAPI staining using Cytation 5 microscope (Agilent BioTek). GraphPad Prism software version 11 was used to calculate IC_50_ values with nonlinear regression analysis with a variable slope from three independent experiments with three replicates in each experiment.

### 2.5 Time of addition effect of compounds on HCoV-OC43 infection

Time of addition assays were performed as described above in 2.3, with following modifications: Cells were treated 2h before infection (-2 h to 0 h); during infection (0 h to 2 h); or after virus adsorption (2 h – 24 h). At 0 h, all cells (except the mock control) were infected with HCoV-OC43 (MOI of 0.1 FFU/cell) for 2 h at 33°C, after incubation virus inoculum was removed and cells were washed with RPMI and replaced with infection media (RPMI containing 2% FBS and 1% PEST). In the 0 h to 2 h time point virus was added together with the compound. Wells were divided up respective to the time of addition and presence of compound of the post infection with 2 h intervals (0 h to 2 h, 2 h to 4 h, 4 h to 6 h, 6 h to 8 h, 8 h to 10 h and 10 h to 24 h) for a total experiment time of 24 h from the point of infection (Figure 6A). At 24 h post infection, cell supernatants were discarded and HCoV-OC43 cellular infection was assessed following antibody staining.

### 2.6 CC_50_ value determination of anthracyclines compounds

HCT-8 cells (approximately 1.8 × 10^4^ cells/well) were seeded in 96-well Nunclon delta surface white plates (Thermo Fisher). Approximately 24 h later, compounds were serially diluted in two-fold steps from 40 to 0.156 μM in RPMI-1640 supplemented with 2% FBS and added to the cells and incubated for 24 h at 37°C in 5% CO_2_. Cell viability was assessed using a CellTiter-Glo Luminescent Cell Viability assay kit (Promega) following the manufacturer instructions. Luminescence was measured using CLARIOstar Plus plate reader (BMG Labtech).

### 2.7 DNA damage determination

HCT-8 cells (approximately 5 × 10^4^ cells/chamber) were seeded on 4-Chamber tissue culture glass slides (BD Biosciences). Approximately 24 h later, compounds were added to a final concentration of 1 µM to the cells and incubated for 16 h at 37°C in 5% CO_2_. The slides were washed and fixed with 4 % PFA for 20 minutes at RT. DNA damage was visualized by DAPI staining and immunofluorescence microscopy using a rabbit anti-pKAP1 antibody (A300-767A, Bethyl) together with a mouse anti-gH2AX antibody (05-636, Millipore) in a 1:100 dilution in blocking buffer (PBS with 2.5 % BSA and 0.25 % Triton) for 1 h. After washing a secondary goat anti-rabbit Alexa Fluor 568 antibody was used together with a donkey anti-mouse Alexa Fluor 488 antibody in a 1:1000 dilution for 1h. Images were taken using a Nikon eclipse 90i microscope equipped with Hamamatsu ORCA-ER camera, Nis element AR 3.2 software and CoolLED pE-300 light source with a 40x Plan Fluor, NA 0.75 objective. Acquisition settings have been kept uniformly to allow comparison of different treatments. Background was subtracted and images processed using FIJI/ImageJ.

### 2.8 Effects of anthracyclines on SARS-CoV-2 infection

A549-ACE2 cells were seeded in 96-well plates at a density of 1.2x10^4^ cells/well in complete Dulbecco’s Modified Eagle’s Medium (DMEM) and incubated for 24 h at 37°C (5% CO_2_). Cells were inoculated with SARS CoV-2 variants (Wuhan, Delta, Omicron BA.2 and XBB.1.9.2.5.1.3) (MOI 0.1) together with compounds at 1 and 2.5 μM concentrations. The plates were incubated for 16 h at 37°C in 5% CO_2_. SARS CoV-2 infection was detected following incubation with primary rabbit anti-SARS-CoV-2 nucleocapsid monoclonal antibody (40143-R001, Sino Biological) and secondary anti-rabbit Alexa Fluor 647 antibody (A-31573, Invitrogen); cellular nuclei were stained with 0.1% DAPI. Virus infection and number of cells in each well were quantified using Cytation 5 microscope (Agilent BioTek).

VeroE6 cells were seeded in 24-well plates at a density of 8x10^4^ cells/well in complete Dulbecco’s Modified Eagle’s Medium (DMEM) and incubated for 24 h at 37°C (5% CO_2_). Cells were inoculated with SARS-CoV-2 strain delta (MOI=0.05) in presence of compounds at 2.5 μM or 20 μM (Merafloxacin) concentrations in DMEM 1% FBS 1% PeST for 48h at 37°C (5% CO_2_). The media were discarded and the cells were lysed using RIPA buffer with 1% protease inhibitor for Western blot analysis.

### 2.9 Generation of DNA templates and *in vitro* transcription

All used plasmids and oligonucleotides are listed in Tables S1 and S2, respectively. Constructs were generated in a similar fashion as described in [11]. Briefly, a DNA template containing a T7 promoter, leader region and Kozak sequence followed by an N-terminal 3x FLAG-tag and linker and the wildtype FSE locus of HCoV-OC43 or SARS-CoV-2 with most of the *nsp12* gene and a *Sma*I restriction site was synthesized by Eurofins Genomics into the vector pEX-A258 (see Figure 3D). By Q5 site-directed mutagenesis PCR we disrupted the slippery sequence of the frameshift context to generate an in-frame control resulting in maximal expression of the -1-frame product (IFC; pEX-HCoV-OC43/SARS-CoV2-IFC) and a stop control resulting in minimal expression of the -1-frame product (Stop; pEX-HCoV-OC43/SARS-CoV2-Stop). *In vitro* transcription was done with the MEGAscript T7 Transcription Kit (Invitrogen) according to the manufacturer’s instructions with 1 µg of *Sma*I digested pEX-constructs as template.

**Figure 3.**
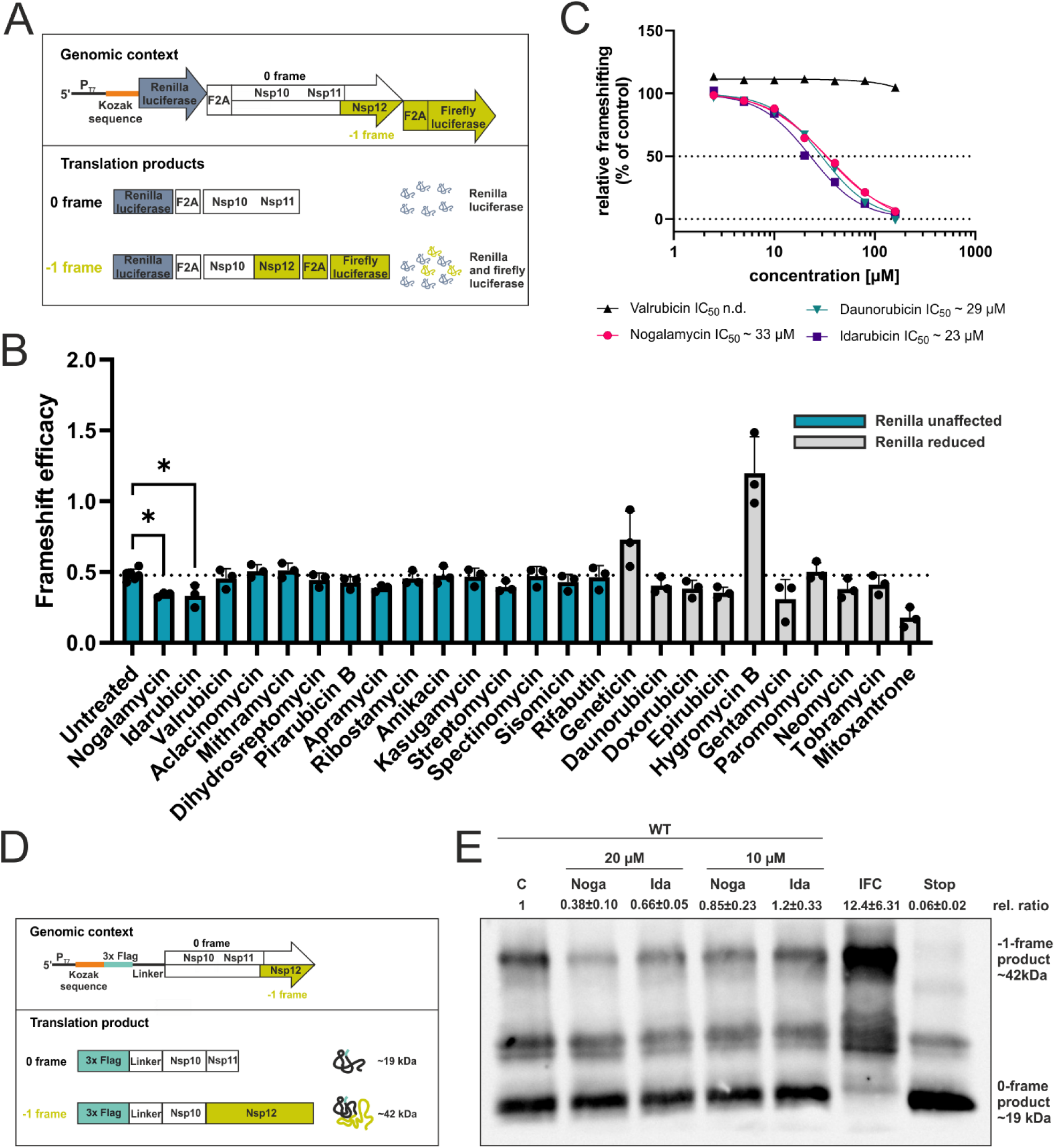
Idarubicin and Nogalamycin reduce – 1 PRF in HCoV-OC43. **(A)** Schematic representation of the dual luciferase reporter gene construct. Firefly luciferase is only expressed upon frameshifting, while *Renilla* luciferase is constitutively expressed. F2A – StopGo signal. **(B)** Nogalamycin and Idarubicin reduce frameshifting during *in vitro* translation. *In vitro* translation of the Dual luciferase-HCoV-OC43-FSE reporter by rabbit reticulocyte lysate. Experiments were carried out three times, except for the untreated control (n=6). Mean and corresponding standard deviation are shown. The *p*-values are indicated as * *p*<0.05. **(C)** Effective compounds show a dose dependent response. Experiments have been performed as described in (B). All values have been normalized, relative to the control translation as 100 %. Experiments were carried out three times. **(D)** Schematic representation of the 3xFlagtag reporter gene construct. **(E)** Nogalamycin and Idarubicin reduce frameshifting during *in vitro* translation. *In vitro* translation of the 3xFlagtag-HCoV-OC43-FSE reporter by rabbit reticulocyte lysate. Experiments were carried out three times. One representative Western blot is shown. Relative ratios have been calculated by quantifying the -1-frame product and 0-frame product intensities using ImageJ and normalization against the control. C – control; IFC- in frame control (maximal -1-frame production); Stop – Stop control (maximal 0-frame production).

To generate Dual-luciferase reporter constructs (see Figure 3A), the FSE locus was amplified by PCR using the 5’-*Xho*I and 3’-*BamH*I primers. After digestion the insert was ligated into the *PspX*I and *Bgl*II digested pSGDlucV3.0 vector (pSGDlucV3.0 was a gift from John Atkins Addgene #119760 [40]) generating pSGDluc-HCoV-OC43/SARS-CoV-2-reporter. After amplification with T7-fw and pSGDLuc_IV_rev, 200 ng of the PCR product were used as template for *in vitro* transcription with the MEGAscript T7 Transcription Kit (Invitrogen) according to the manufacturer’s instructions.

### 2.10 *In vitro* translation

*In vitro* translation was carried out with the nuclease-treated rabbit reticulocyte lysate Kit (Promega) according to the manufacturer’s instruction. Briefly, 1.2 µg of *in vitro* transcribed RNA was used in a 25 µl standard reaction at 30°C for 90 minutes in presence or absence of the tested compound. Total DMSO concentration in the reaction mixture was 1 %. Translation products were further used for SDS-PAGE analysis (pEX-3xFlag constructs) or Luciferase activity measurements (pSGDLuc constructs).

### 2.11 SDS-PAGE and Western blot analysis

Of the *in vitro* translation product, 5 µl were mixed with 20 µl 1x SDS sample buffer (2% SDS, 0.1% bromophenol blue, 1% 2-mercaptoethanol, 25% glycerol, 50 mM Tris/HCl, pH 6.8) and boiled for 10 min at 95°C. Afterwards 7-10 µl of the supernatant was loaded and separated by SDS gel electrophoresis in 5% stacking and 12% separating gels. For the virus infected cell lysates, total protein concentration was determined via the Pierce BCA Protein assay kit (Thermo scientific) and 15 µg protein (for NSP8 and NSP12 detection) or 5 µg protein (for Nucleocapsid and Actin detection) were mixed with 6x SDS sample buffer boiled for 10 min at 95°C and separated by SDS gel electrophoresis in 5% stacking and 10% separating gels.

By semidry blotting, the proteins were transferred onto a nitrocellulose membrane (BioTrace NT, Pall Corporation) and a mouse anti-Flag antibody (Sigma-Aldrich; F3165) was used in a 1:2000 dilution, The next day, the membranes were washed 3 times with TBST and incubated with an anti-mouse-HRP conjugate antibody (A9044, Sigma-Aldrich) in a 1:10000 dilution. For SARS-CoV-2 protein detection a rabbit anti-SARS-CoV-2 nucleocapsid (40143-R001, Sino Biological) was used in a 1:8000 dilution, a mouse anti-NSP8 antibody (MAB12502, R&D Systems) in a 1:5000 dilution, anti-NSP12 (9267, ProSci Inc.) in a 1:500 dilution, rabbit and anti α-actin (A2066, Sigma-Aldrich) in a 1:7500 dilution, respectively. Chemiluminescence signals were detected by incubating membranes with Amersham ECL prime western blotting detection reagent (Cytiva) for 3 minutes. For quantification, band intensities were measured with ImageJ.

### 2.12 Dual luciferase activity assay

Luciferase activity was measured with the Dual-Glo Luciferase Assay system (Promega). Briefly, 20 µl of the *in vitro* translation product was transferred into a white 96-well plate and mixed with 100 µl of Dual-Glo Luciferase reagent. After 10 minutes of incubation at room temperature, firefly luciferase activity was measured in a multimode microplate reader (Tecan, Spark). Afterwards 100 µl of freshly prepared Dual-Glo Stop & Glo reagent is added and *Renilla* luciferase activity is measured after another 10-minute incubation at room temperature. To calculate frameshift efficiency first, the relative frameshift ratio for all samples was calculated (firefly/*Renilla* activity). Next the ratio for the stop control was subtracted from all other samples (sample ratio – stop control ratio). Lasty, the resulting ratio is divided by the ratio of the in-frame control (sample-stop ratio/IFC-stop ratio) resulting in a normalized ratio or relative response ratio

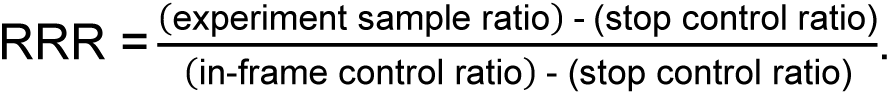

#### DMS modification of *in vitro* transcribed RNA

For DMS treatment HCoV-OC43 FSE RNA was generated from a T7-promoter containing DNA template (with T7-fw-fullFSE or T7-slippery-fw and fullFSE-iv-rev primers). The RNA was treated by DMS and library generation was performed as described in [41,42,12]. For this, 10 µg of *in vitro* transcribed RNA in 10 µl H_2_O, was denaturated at 95°C for 3 minutes and quickly put on ice. 88 µl of DMS folding buffer (300 mM sodium cacodylate with 6 mM MgCl_2_, [43]) were added and the RNA was allowed to refold for 5 minutes at 37°C. Of the compounds 1 µl (2 mM) was added, resulting in a final concentration of 20 µM and incubated for another 5 minutes. Then the RNA was treated with 2 µl DMS for 3 minutes at 37°C while shaking. The reaction was stopped by adding 60 µl β-mercaptoethanol and the RNA precipitated with ethanol.

For DMS modification with the compound present during folding following modifications were applied to the protocol above. For this 2 µM *in vitro* transcribed RNA in 10 mM MOPS-Na pH 7 and 1 mM EDTA was denatured at 90°C for 3 minutes and quickly put on ice. Refolding of the RNA was initiated by adding 5 µl of the denatured RNA to 45 µl of pre-warmed solution that yields a final concentration of components: 0.2 µM RNA, 20 µM compound of interest, 1 % vol/vol DMSO, 300 mM Na-cacodylate pH 7 and 10 mM MgCl_2_. The RNA was allowed to fold for 20 minutes at 50 °C then cooled down to room temperature for 3 minutes. DMS treatment was performed for 6 minutes at 37 °C by adding 5.6 µl of 5 % vol/vol DMS solution in ethanol. The reaction was stopped by quenching the DMS with an equal volume (56 µl) of 2-mercaptoethanol. The RNA was recovered by an overnight precipitation with ethanol using glycogen as co-precipitant.

### 2.13 Ligation-based DMS-MaPseq library generation

After precipitation, the FSE-RNA was used for library preparation as described in [41]. For this the RNA was prepared for adapter ligation by dephosphorylation of the 5’ end with 5’ Polyphosphatase (Biosearch Technologies) and cleaned up with the RNeasy MinElute cleanup kit (Qiagen). Next the 5’ RNA adapter was ligated to the FSE-RNA with T4 RNA Ligase 1 (New England Biolabs) and afterwards gel purified from a 6 % denaturing polyacrylamide gel by excision and ethanol precipitation, to remove non-ligated adapters. Next, the 3’ ends of the RNA were dephosphorylated with T4 Polynucleotide Kinase (Thermo scientific) and cleaned up with the RNeasy MinElute cleanup kit. Now the adenylated 3’DNA adapter was ligated to the RNA with T4 RNA Ligase 2, truncated (New England Biolabs) and afterwards ethanol precipitated and cleaned up with the AMPure XP beads (Beckman Coulter) with a 1.6 x ratio of magnetic beads to volume to remove non-ligated adapters. The cDNA was synthesized with Induro Reverse Transcriptase (New England Biolabs). The RNA was concentrated to a volume of 4.5 µl by a SpeedVac concentrator. The RNA was mixed with 1 µl of 10 mM dNTPs, 2 µl of 10 µM RT-primer and 2.5 µl H_2_O and denatured for 5 min at 65°C after cooling down on ice. Afterwards, 4 µl of 5x Induro RT buffer, 0.2 µl Ribolock (Thermo scientific) 1 µl Induro RT and 4.8 µl H_2_O were added and the cDNA was synthesized for 10 minutes at 55 °C with a 1-minute inactivation at 95°C. To degrade the RNA 1 µl of 5 M NaOH was added and incubated at 95°C for 3 min. The first strands of cDNA were ethanol precipitated and used for amplification and addition of Illumina Trueseq Indexes with Phusion Mastermix (Thermo scientific) for 12-15 cycles. Afterwards the PCR products were enriched by a second round of PCR with a maximum of 10 cycles and purified with QIAquick PCR purification Kit (Qiagen) and further cleaned up with the AMPure XP beads in a 1:1 ratio. The resulting libraries were sequenced on an Illumina MiSeq instrument using either 2 x 75 (V3 chemistry or 2 x 150 (v2 chemistry) paired-end sequencing.

### 2.14 Mapping, quantification of mutations and computing

The sequencing results where further processed using DREEM [44] as previously described [43]. Briefly, FASTQ files were trimmed by TrimGalore, if adapter sequences have not been removed before already. The reads were mapped to the reference sequence with Bowtie2 [45], Sam files converted into BAM files using Picard Tools SamFormatConverter and bit vectors and corresponding DMS reactivities generated by DREEM.

### 2.15 Normalization and folding of the FSE according to the DMS reactivity

DMS reactivities were normalized to a scale of 0 – 1 according to [43]. First a median among the top 5% of DMS reactivities was calculated. The DMS reactivities were normalized to this value and values greater than 1.0 were winsorized by setting them to 1.0 to still include them in the analysis. The FSE-RNA was folded by the ShapeKnots algorithm [46] of the RNAstructure package [47] with the parameters -m3 to generate three structures and -dms to use the normalized mutations rates as folding constraints. Connectivity table files were converted to dot-bracket format by ct2dot from the RNAstructure package.

### 2.16 Fluorescence-quenching assay

Anthracycline and RNA interaction was investigated by utilizing the intrinsic fluorescence of many anthracyclines which could be quenched by direct RNA-binding as has been shown for Idarubicin-DNA interactions [48] and Idarubicin-RNA interactions [36]. Briefly, 90 µl of *in vitro* transcribed RNA were heated to 65°C for 2 min and cooled to allow RNA secondary structure formation. The RNA was transferred into a black 96-well plate and 10 µl of the anthracyclines, dissolved in DMSO were added to a concentration of 4 µM and incubated for 1 h at room temperature. The fluorescence intensities were recorded using a multimode microplate reader (Tecan, Spark) at the excitation wavelength 482 nm and emission wavelength of 571 nm. Additionally, an emission spectrum was recorded at the excitation wavelength 482 nm from 520 to 850 nm with a 10 nm measurement interval.

## 3. Results

### 3.1 Idarubicin and Nogalamycin reduce -1 PRF in HCoV-OC43

Owing their RNA-binding properties, we hypothesized that aminoglycosides and/or anthracyclines could inhibit coronavirus replication by affecting the frequency of - 1 PRF. To test this, we first examined whether these compounds influence the frameshift efficacy *in vitro* by testing all commercially available aminoglycosides and anthracyclines (n=25). First, we chose to examine the ability of the compounds to affect -1 PRF using an *in vitro* translation system as described in [11], where the FSE of HCoV-OC43 was inserted in a dual luciferase system flanked with StopGo (F2A sequences), generating multiple separate proteins instead of one large fusion protein [49,50] (Figure 3A).

In this reporter system, the *Renilla* luciferase is continuously expressed, whereas the firefly luciferase is only synthesized upon -1 frameshifting. By monitoring the ratio of firefly versus *Renilla* luciferase expression, the frameshifting efficiency can be determined. An in-frame control (IFC), where the firefly luciferase is in the same frame as the *Renilla* luciferase, was used as a positive control (maximal firefly expression) and a construct harbouring a stop codon in the slippery sequence was used as a negative control (0-frame, no firefly expression). Using this system, a frameshift ratio of ∼ 50 % was observed, fitting to previously reported values for other coronaviruses (25-75 % [11,25,51–53]) (Figure 3B). When examining the frameshifting efficiency in presence of compounds, we observed a significant reduction by Idarubicin and Nogalamycin. Several of the compounds (n=10) affected general translation (Figure 3B, grey bars and Figure S1). Among those was Geneticin, an aminoglycoside, previously reported to inhibit -1 PRF in SARS-CoV-2 infection [16]. For IC_50_ determination, we thus examined the efficacy of the most prominent anthracyclines (Idarubicin, Nogalamycin and Daunorubicin), on -1 PRF and included Valrubicin as a negative control (Figure 3C). The largest effect was observed using Idarubicin, followed by Nogalamycin and Daunorubicin. The concentrations required to inhibit 50% of the frameshifting (IC_50_) was 20.5 µM for Idarubicin; 35.4 µM for Nogalamycin and 36.4 µM for Daunorubicin, respectively, whereas Valrubicin did not affect -1 PRF.

To further verify our data and exclude possible interference of the anthracyclines on luciferase activity, an additional *in vitro* translation assay was conducted where the FSE was inserted into a Flag-tag reporter system (Figure 3D). This assay generates a short translation product (∼19 kDa) in absence of frameshifting whereas a longer translation product (∼42 kDa) is produced upon frameshifting. Using this system in presence of Nogalamycin and Idarubicin showed a similar reduction in frameshifting as observed using the dual luciferase assay, confirming an effect on -1 frameshifting (Figure 3E).

Next, we asked whether the anthracyclines could have a synergistic effect on -1 PRF. We employed the *in vitro* translation dual luciferase system and used equal amounts of Idarubicin and Nogalamycin. No synergistic effect of the anthracyclines was observed even with 10 µM of both Idarubicin and Nogalamycin, respectively, indicating that the compounds act by a similar mechanism of action and/or have the same binding pocket in the FSE (Figure S2).

### 3.2 Anthracyclines bind to the HCoV-OC43 FSE

Next, we were interested to examine whether the anthracyclines interact directly with the FSE. To answer this, we first investigated if the RNA structure of the FSE was altered upon binding the drugs. For this, *in vitro* transcribed FSE was refolded in presence of the anthracyclines and subjected to dimethyl sulfate mutational profiling with sequencing (DMS-MaPseq [54]) where non-interacting adenines and cytosines can be distinguished from bases binding another base or a molecule (Figure 4)

**Figure 4.**
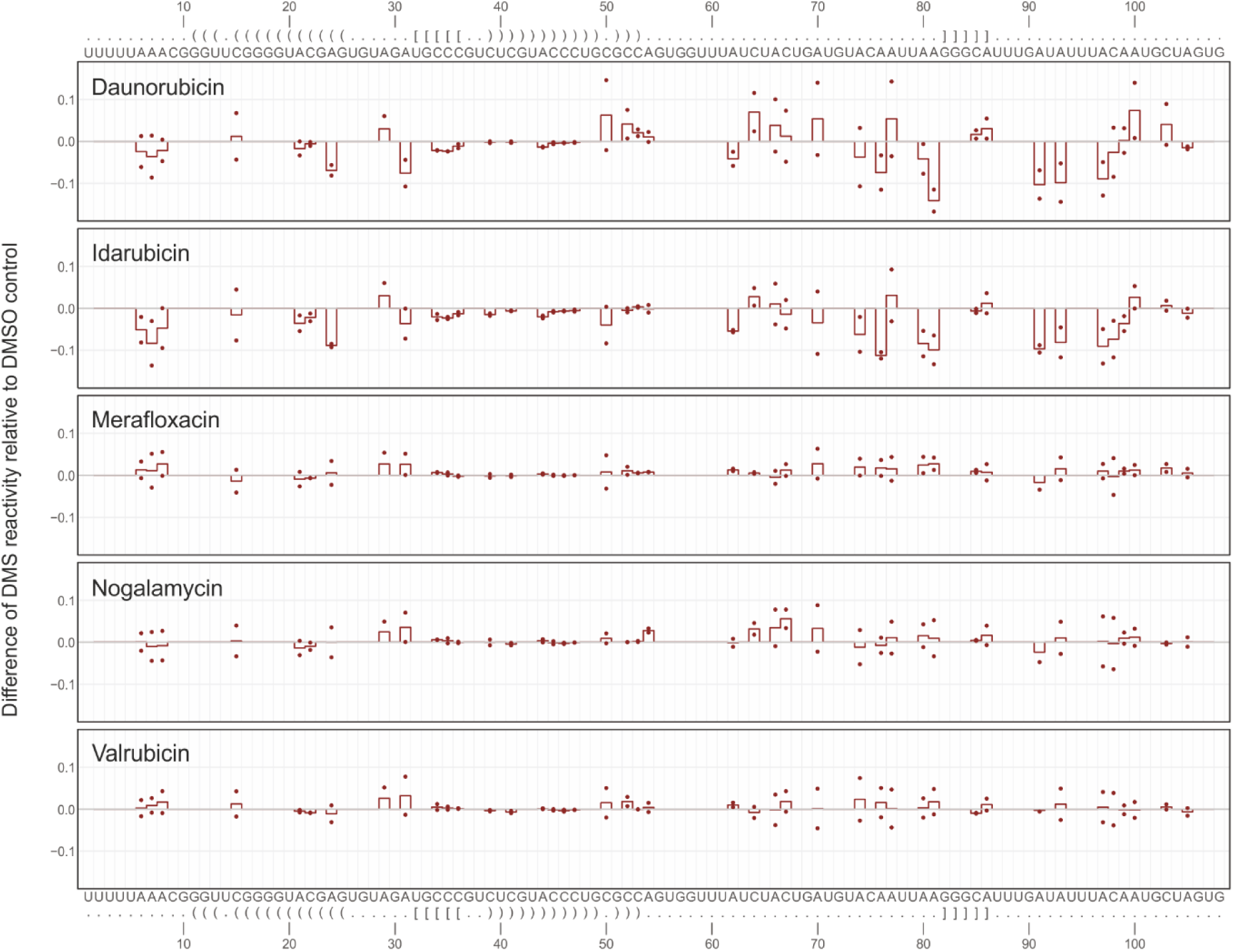
DMS-reactivity is altered when Daunorubicin and Idarubicin are added during folding of the FSE. Normalized DMS reactivities in absence or presence of compound relative to DMSO control. Bases are shown above and below plot. Negative values indicate that the base interacts with another base or with a molecule.

The profiling showed that the basic pseudoknot structure of the frameshift element was still maintained, also in the presence of anthracyclines able to reduce -1 PRF (Figure 4). However, Daunorubicin and Idarubicin reduced DMS-reactivity, particularly at the positions A76, A80 and A81 compared to DMSO (Figure 4). Interestingly, these bases are located in a loop between stem 2 and stem 3, which could accommodate molecule binding (Figure 1B). We did not observe any changes of the DMS-reactivity in presence of Nogalamycin, Merafloxacin or Valrubicin as compared to the DMSO control.

Allowing the FSE to fold before adding the anthracyclines did not result in differences in altered DMS-reactivity, (Figure S3) indicating that the compounds need to be present with the FSE during folding for a stable interaction.

To further examine the anthracycline-FSE interactions, we took advantage of the intrinsic fluorescence that many anthracyclines exhibit and that can be quenched by direct RNA-binding as has been shown for anthracycline-DNA interactions [48]. In line with a direct interaction between the anthracyclines and RNA, Idarubicin, Nogalamycin as well as Daunorubicin exhibited strong intrinsic fluorescence that was dramatically quenched in the presence of the FSE RNA (Figure 5).

**Figure 5.**
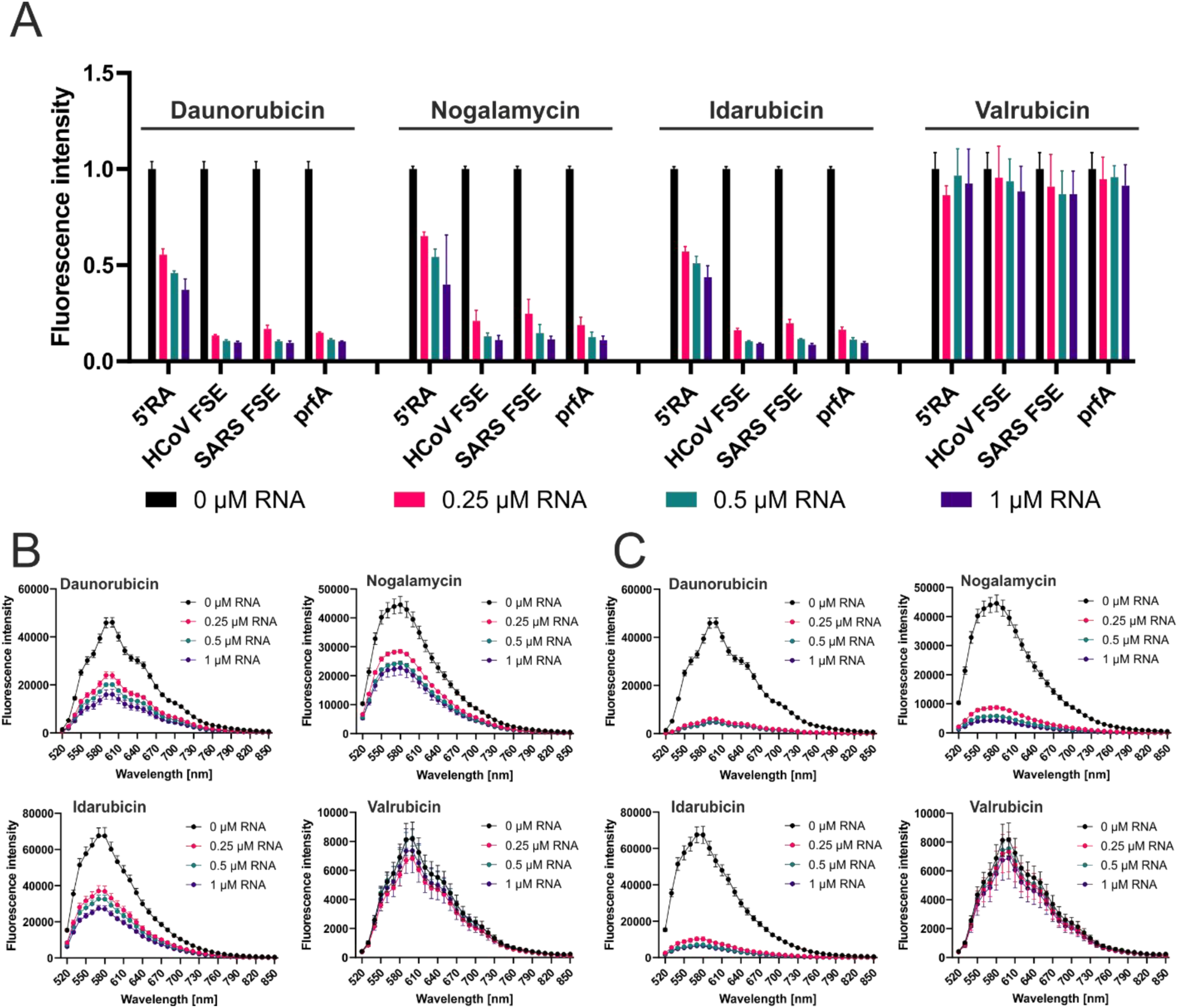
Intrinsic anthracycline fluorescence is quenched in presence of HCoV-OC43-FSE RNA. **(A)** Fluorescence intensities were measured at 482 nm excitation and 571 nm emission in absence or presence of indicated concentrations of RNA. Experiments were carried out three times. Mean and corresponding standard deviation are shown. Emission spectra recorded after excitation at 482 nm of the different compounds in absence and presence of 5’RA RNA **(B)** and HCoV-OC43 RNA **(C)**.

Valrubicin, which was unable to inhibit -1 PRF, showed no quenching by the presence of RNA at any concentration, strongly indicating that Valrubicin does not bind the FSE, thus corroborating the data from the frameshift and the DMS-MaPseq experiments (Figure 3, 4 and S3). To examine specificity of the anthracyclines, we measured the level of quenching in presence of other RNA molecules. An unrelated, unstructured RNA, the 5’ RNA adapter (5’RA), previously used for library preparation for DMS-MaPseq (Table S2) was tested as well. The potential of this unstructured RNA to quench the anthracyclines was much lower compared to the HCoV-OC43 FSE. Additionally, we tested the SARS-CoV-2 FSE, as well as a bacterial RNA thermometer hairpin structure [55]. In line with the abilities of anthracyclines to bind structured nucleic acids [30–32,36], we observed binding to a similar extent as the HCoV-OC43 FSE (Figure 5) for these structures.

### 3.3 Anthracyclines reduce HCoV-OC43 infection

Given the fact that some anthracyclines interacted with the HCoV-OC43 FSE and modulated frameshifting efficiency, we next asked whether the effect of the anthracyclines on -1 PRF could reduce the ability of HCoV-OC43 to cause infection. To examine this, we analysed HCoV-OC43 infection of HCT-8 cells in presence of compounds using immunofluorescence microscopy (Figure 6).

All previously tested (Figure 3) anthracyclines and aminoglycosides (n=25) were re-tested for antiviral activity, and a prominent antiviral effect was only observed for a subset of anthracyclines. Most strikingly, Idarubicin reduced infection by ∼40 % compared to the control already at 1 µM (Figure 6A) and almost completely inhibited infection (∼90 % reduction) at 5 µM (Figure 6B and C). Other anthracyclines (Nogalamycin, Daunorubicin, Doxorubicin and Epirubicin, respectively) also reduced infection effectively at 5 µM by ∼30-50 %, whereas aminoglycosides did not. Interestingly, although it has been reported to inhibit SARS-CoV-2 entry, we did not observe an effect by Mitoxantrone, an anthraquinone with a similar molecular structure to Daunorubicin [56]. Due to their effect on both -1 PRF and cell infection, the concentrations that inhibit 50% of the infection (IC_50_) were determined for Idarubicin, Nogalamycin and Daunorubicin (Figure 6D). Valrubicin was included as negative control as it did not affect -1 PRF nor showed an antiviral effect at low concentrations (Figure 3 and Figure 5). The IC_50_ values for Idarubicin, Daunorubicin and Nogalamycin, were 2.8 µM; 5.8 µM and 6.2 µM, respectively, whereas Valrubicin only reduced infection at higher concentrations (IC_50_ 15.5 µM, Figure 6D).

To further investigate the mechanism of action, we conducted time of addition experiments to further determine at what point the anthracyclines exhibit their antiviral effect. For this, HCT-8 cells were treated with Nogalamycin or Idarubicin for 2 h intervals before, during and after infection with HCoV-OC43 (Figure 7A).

**Figure 6.**
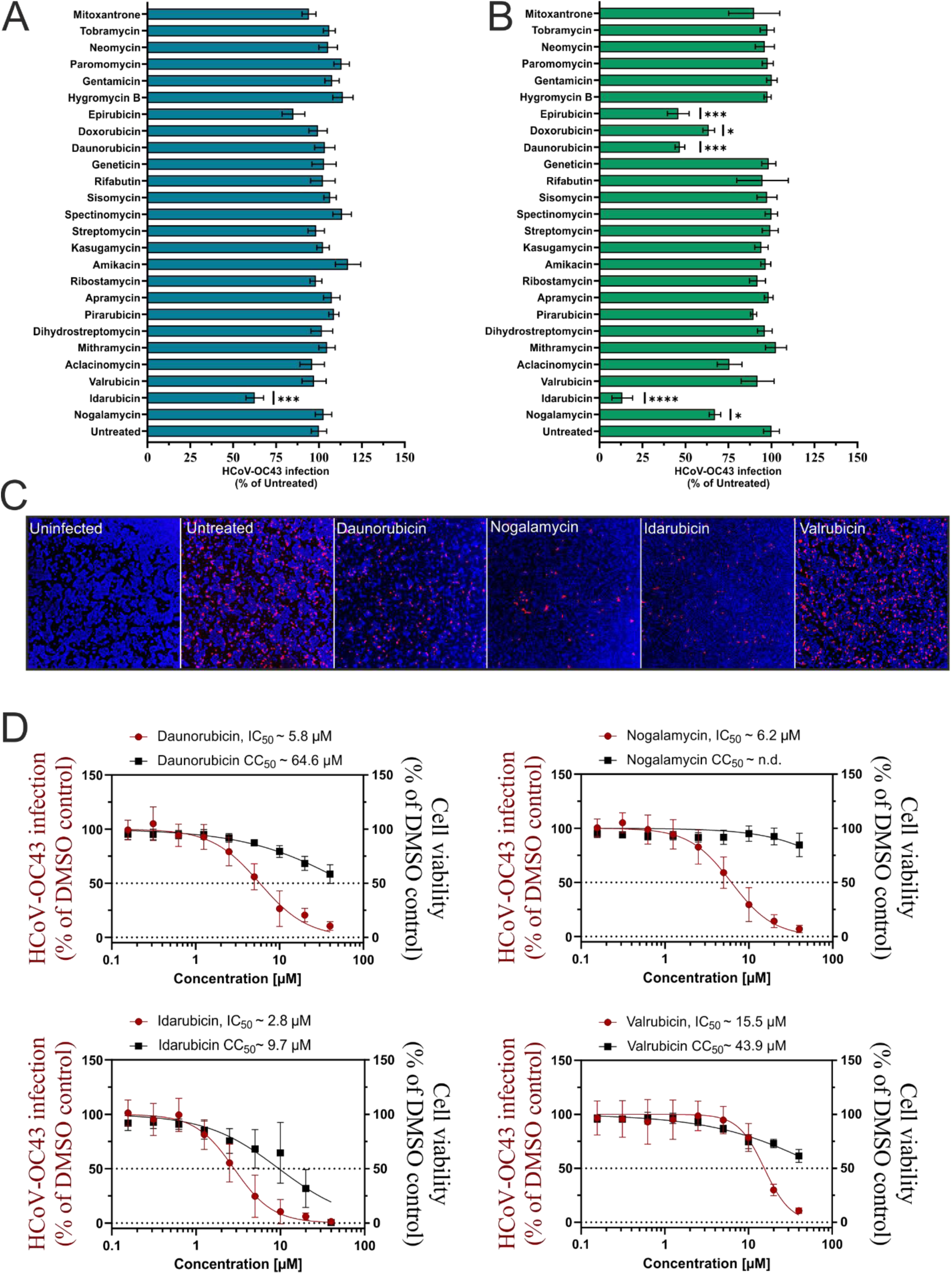
Anthracyclines reduce HCoV-OC43 infection and replication. Inhibition of HCoV-OC43 infection by aminoglycosides and anthracyclines. HCT8 cells were infected with HCoV-OC43 (MOI =0.1) for 16h in presence of 1 **(A)** and 5 µM **(B)** of the indicated compound. Infection was quantified by immunofluorescence with a nucleocapsid antibody and a secondary Alexa Fluor 647 antibody. Results were generated with three independent experiments, each with three replicates. Mean and corresponding standard deviation are shown. The *p*-values are indicated as * *p*<0.05; \*\*\**p*<0.001; \*\*\*\**p*<0.0001 **(C)** One representative microscopy picture at 5 µM used to generate data shown in **(B)**. Cell nuclei were stained with 0.1 % DAPI. **(D)** Effective compounds show a dose dependent response. Experiments were performed as described in **(A)**. Cell viability was determined by Cell Titer Glow Cell Viability Assay Kit (Promega). Results were generated with three independent experiments, each with three replicates.

**Figure 7.**
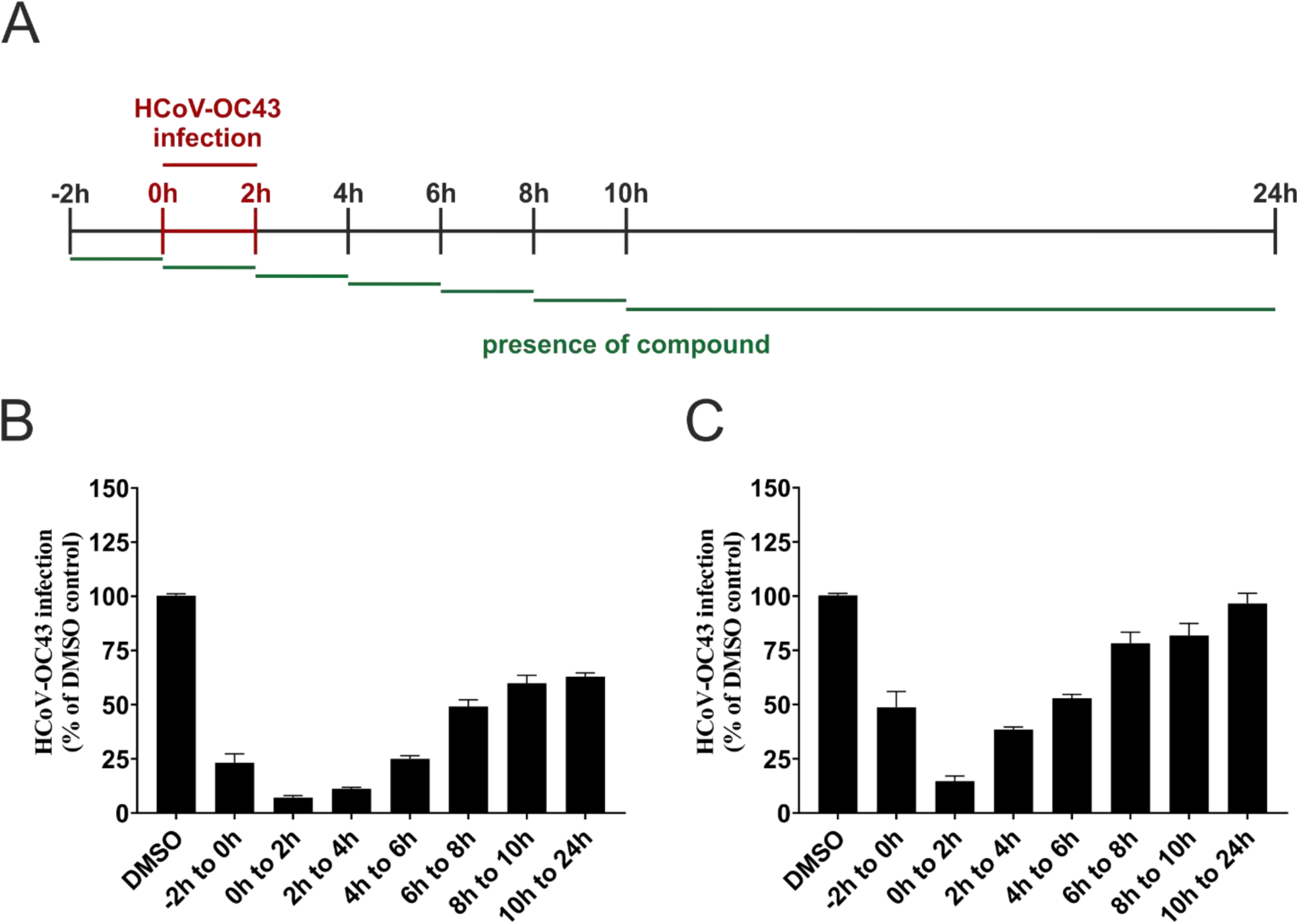
Nogalamycin and Idarubicin primarily act during early stages of infection. **(A)** Time of addition study, indicating the presence of compound at different stages. For time of addition studies HCT8 cells were infected with HCoV-OC43 (MOI= 0.1, t= 0h for 2 h) and incubated for up to 24 h after treatment with 10 µM Idarubicin **(B)** or Nogalamycin **(C)** at different timepoints for 2 h each or 14 h. After 2 h compounds and virus inoculum (t= 2h) were removed, cells were washed and further incubated until evaluation. The viral infection was determined by immunofluorescence with a nucleocapsid antibody and a secondary Alexa Fluor 647 antibody. Results were generated with four independent experiments, each with three replicates. Mean and corresponding standard deviation are shown.

After 24 hours, the viral infection was quantified as before (Figure 6A). Our results indicate that Idarubicin and Nogalamycin acted during the early stages of infection, as we observed a strong reduction of infection (up to 90%) if the anthracyclines were added early during infection (0h to 2h), thus affecting entry, initial translation and replication (Figure 7B and C). The effect was stronger for Idarubicin than for Nogalamycin, with Idarubicin showing an antiviral effect if added also later during the infection (Figure 7A and B).

### 3.4 Potent anthracyclines cause DNA damage

Being anti-cancer drugs, anthracyclines generally show strong effects on cell growth, where Idarubicin and Daunorubicin inhibit the progression of DNA topoisomerase II by intercalating with DNA, thereby affecting transcription and replication and eventually causing double strand breaks, [57–59]. Considering this, we tested the cytotoxicity of the drugs. Daunorubicin and Valrubicin did only affect cell viability at relatively high concentrations (CC_50_ values of ∼ 64.6 µM and ∼43.9 µM, respectively) whereas Idarubicin affected cell viability at lower concentrations (CC_50_ value at ∼ 9.7 µM). Nogalamycin did not affect cell viability.

Although the anthracyclines do not kill cells at lower concentrations, the concentrations could still be sufficient to affect cell physiology and have long-term consequences. The cell infection inhibition could thus be hypothesized to be mediated by indirect effects on cell physiology. To answer this, we examined whether inhibition of viral infection by anthracyclines could be explained by effects on cell-toxicity by DNA damage. HCT-8 cells were grown on microscopy glass slides in presence of 1 µM compound. Using immunofluorescence we investigated the presence of early DNA damage markers; γH2AX, which is the Ser-139 phosphorylated version of the minor histone H2A variant [60,61] and pKAP1, the phosphorylated KRAB associated protein 1 [62,63]. As a positive control we used the well-known topoisomerase II poison, Etoposide [64,65]. While the DMSO control only showed minor DNA damage, we observed a strong signal for the pKAP1 and γH2AX markers among all tested compounds at low concentrations (Figure 8).

**Figure 8.**
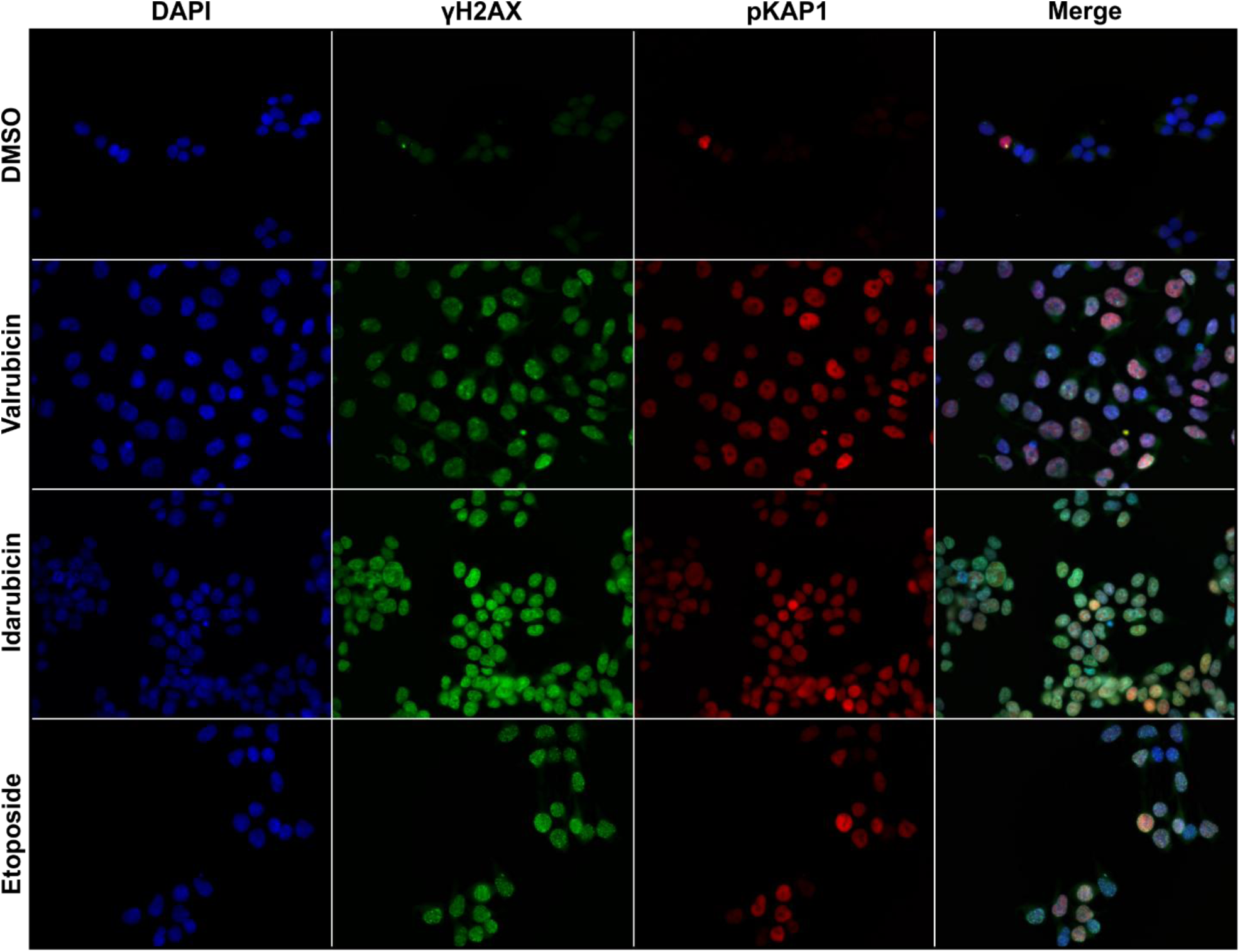
Idarubicin and Valrubicin cause DNA damage. HCT8 cells were grown on glass slides and treated with 1 µM compound or DMSO for 16 h, before fixation and staining with DAPI (blue). The indicated primary antibodies detect early DNA damage markers γH2AX (green) and pKAP1 (red). Experiments were carried out three times. Images have been acquired with the same settings to allow comparison. One set of representative images is shown.

Interestingly we also detected strong signals for our negative control, Valrubicin, suggesting that DNA damage alone cannot be the reason for the reduced infection during HCoV-OC43 infection (Figure 6).

### 3.5 Anthracyclines can reduce SARS-CoV-2 infection but do not inhibit -1 PRF

Since we observed binding of our potent anthracyclines to the SARS-CoV-2 FSE (Figure 5A) which is structurally similar to the HCoV-OC43 FSE (Figure 1B), we were interested in examining whether these compounds could also affect SARS-CoV-2 infection. Different strains of SARS-CoV-2 were used to infect A549-ACE2 cells in the presence of our chosen compounds. Interestingly, all tested compounds, except Daunorubicin significantly reduced SARS-CoV-2 infection, with Nogalamycin showing the strongest effect (Figure 9A and Figure S4). Nogalamycin reduced Delta variant infection by ∼ 50 % already at 1 µM (Figure 9A). A similar trend was observed for the other SARS-CoV-2 strains as well (Figure S4).

**Figure 9.**
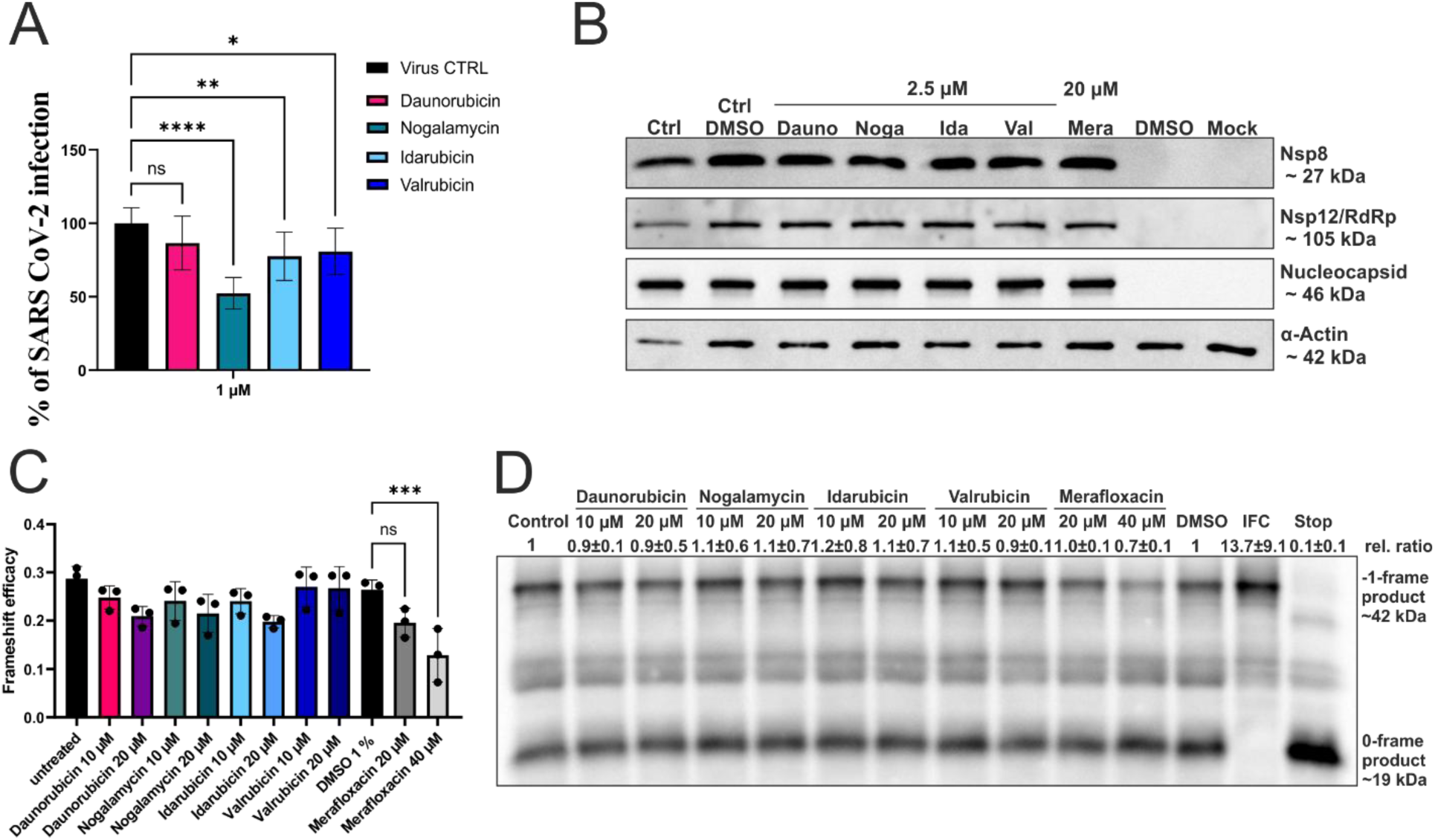
Anthracyclines reduce SARS-CoV-2 infection but do not affect -1 PRF. **(A)** Inhibition of SARS-CoV-2 strain delta infection by anthracyclines. A549-ACE2 cells were infected with SARS-CoV-2 (MOI = 0.1) for 16h in presence of 1 µM of the indicated compound. The viral infection was determined by immunofluorescence with a nucleocapsid antibody. Results were generated with three independent experiments, each with three replicates. Mean and corresponding standard deviation of all replicates are shown. p values are indicated as \**p*<0.05; \*\**p*<0.01; \*\*\*\**p*<0.0001; ns, not significant. **(B)** Anthracyclines do not inhibit frameshifting of SARS-CoV-2 strain delta during infection. VeroE6 cells were infected with SARS-CoV-2 (MOI =0.05) for 48h in presence of the indicated compound. Ctrl – virus infected control. Ctrl DMSO – virus infected control treated with DMSO. Dauno – Daunorubicin treated infected cells. Noga – Nogalamycin treated infected cells. Ida – Idarubicin treated infected cells. Val – Valrubicin treated infected cells. Mera – Merafloxacin treated infected cells. DMSO – uninfected, DMSO treated cells. Mock – uninfected and untreated cells. **(C)** The anthracyclines show a non-significant effect on frameshifting during *in vitro* translation. *In vitro* translation of the Dual luciferase-HCoV-OC43-FSE reporter by rabbit reticulocyte lysate. Experiments were carried out three times. Mean and corresponding standard deviation are shown. The only significant change was between DMSO and the Merafloxacin (40 µM) samples. \*\*\**p*<0.001), all other changes were non-significant. **(D)** Only Merafloxacin reduces frameshifting during *in vitro* translation. *In vitro* translation of the 3xFlagtag-SARS-CoV-2-FSE reporter by rabbit reticulocyte lysate. Experiments were carried three times. One representative Western blot is shown. Relative ratios have been calculated by quantifying the -1-frame product and 0-frame product intensities using ImageJ and normalization against the control. IFC- in frame control (maximal -1 frame production); Stop – Stop control (maximal 0 frame production).

Next, we examined the effect of the anthracyclines on -1 PRF during SARS-CoV-2 infection (Figure 9B). To determine the putative effect of anthracyclines on -1 PRF during an infection, expression of a protein (NSP8) lying upstream and expression of a protein (NSP12, RdRP) lying downstream of the frameshift element, was examined (see Figure 1A). If the anthracyclines would affect -1 PRF, a decreased NSP12/RdRP expression should be observed, whereas expression of NSP8 should remain unaltered. Vero-E6 cells were infected with the SARS-CoV-2 Delta strain in a similar fashion as described for the frameshift inhibitor Merafloxacin [15]. After 48 h of infection in presence of either Idarubicin, Nogalamycin, Daunorubicin or Valrubicin, respectively, the cells were harvested and protein expression determined (Figure 9B). No effect of the anthracyclines on RdRP-expression was observed, indicating that the compounds did not target -1 PRF of SARS-CoV-2.

To further examine whether the anthracyclines could affect the -1 PRF of SARS-CoV-2, we employed our *in vitro* approaches from the HCoV-OC43 verification (Figure 3A and 3D). We introduced the SARS-CoV-2 FSE into the luciferase and into the Flag-tag *in vitro* translation constructs and examined the effect of the anthracyclines *in vitro* with Merafloxacin included as positive control (Figure 9C and 9D)[15]. As a difference to the effect observed for HCoV-OC43, the anthracyclines did not significantly affect -1 PRF in SARS-CoV-2 neither using the luciferase nor the Flag-tag *in vitro* translation systems (compare Figure 3BE with 9CD). In contrast, Merafloxacin could significantly reduce frameshifting (9CD).

## 4. Discussion

In this study, we have analysed the putative ability of all commercially available aminoglycosides and anthracyclines to affect HCoV-OC43 infection as well as - 1 Programmed Ribosomal Frameshift (-1 PRF). We observed that a subset of anthracyclines (Idarubicin, Nogalamycin and Daunorubicin) significantly affected -1 PRF and bound to the frameshifting stimulating element (FSE) of HCoV-OC43 with some specificity (Figure 3,4 and 5). The same anthracyclines also showed an antiviral effect, being able to reduce HCoV-OC43 infection at low µM concentrations (Figure 6) although anthracyclines also showed a DNA-damaging effect (Figure 8).

Interestingly, despite affecting -1 PRF in HCoV-OC43, none of the tested compounds significantly affected -1 PRF in SARS-CoV-2 (Figure 3BE and 9CD), although they interact with the SARS-CoV-2 FSE equally well as they interacted with the FSE of HCoV-OC43 (Figure 5A). Most likely, sequence and/or structural differences in the FSE elements cause the differences in activities of the anthracyclines (Figure 1B). The most potent anthracycline, Idarubicin, can reduce the ability of HCoV-OC43 to cause infection, reduce -1 PRF and interact with the HCoV-OC43 FSE. Through DMS-MaPseq analysis, we observed reduced reactivity of A76, A80 and A81, when Idarubicin was added during refolding of the FSE RNA, suggesting that Idarubicin could interact with these bases (Figure 4). The bases are predicted to be located at a loop region located between stem 2 and stem 3 (Figure 1B). Interestingly, we did not observe a reduction in the DMS-reactivity when Idarubicin was added after refolding, indicating that the anthracycline interacts with the FSE during transcription/folding of the viral RNA (Figure S3). It has been shown by others that the FSE can fold into a different structure in its full-length form, due to long range RNA interactions [43,66]. Therefore, we cannot exclude a different effect of the anthracyclines on the FSE in a cellular context during infection, when also *de novo* transcription occurs. Further studies are required to examine exactly how Idarubicin interacts with the HCoV-OC43 FSE.

Previous studies have described an effect of the aminoglycoside Geneticin on SARS-CoV-2 infection and -1 frameshifting [16]. In our study, Geneticin did not affect HCoV-OC43 infection (Figure 6A and B). Interestingly, general translation was inhibited *in vitro* in presence of Geneticin at 10 µM (Figure S1). Such indirect translational effects of Geneticin were not detected in the previous study, possibly because the experiments were conducted in cells [16]. For future studies, it is therefore important to examine the effect of new RNA-acting drugs by *in vitro* translation, to ensure correct target determination and avoid possible indirect effects. Although reducing SARS-CoV-2 infection, Mitoxantrone is unable to block HCoV-OC43 infections (Figure 6A), possibly because Mitoxantrone modulates the heparan sulfate-spike complex [56], which is needed by SARS-CoV-2 to enter the cells, but not by HCoV-OC43. Mitoxantrone, which has a similar chemical structure as the anthracyclines affect general translation (Figure S5) and also exhibits topoisomerase II toxic activity and intercalates with DNA [67]. This suggests that the antiviral effect observed for Idarubicin, Daunorubicin, and Nogalamycin is not mediated by indirect DNA-damaging effects. This notion is further corroborated by Valrubicin [68,69], which has a prominent effect on cell toxicity without affecting infection or interacting with the -1 FSE.

The time of addition experiments shed further light onto the potential mechanism by which the anthracyclines act on viral infection. An effect for Idarubicin and Nogalamycin was only observed in the early stages of infection (Figure 7). Addition of compounds between 0 and 2 hours after infection resulted in a strong reduction in viral amount. In MRC-5 cells, where HCoV-OC43 replication is faster than in HCT-8 cells, viral absorption and initial translation and transcription events take place just after infection (T=0h) [70] [71]. In the MRC-5 cell-line, virion release is detected around 9 h after infection [70]. As Idarubicin and Nogalamycin show a much weaker effect on viral inhibition after this timepoint, it suggests that viral egress is not affected (Figure 7BC). It is instead more likely that Daunorubicin, Nogalamycin and Idarubicin affect initial translation and early transcription by reducing -1 PRF. In line with this, other studies have shown that small reductions of -1 PRF, similar to what we observe (∼ 30 % Figure 3B), can significantly reduce infection [12,15,27].

Previous studies have shown that anthracyclines can be potent antiviral compounds not only against SARS-CoV-2 [35] but also Hepatitis B virus [37] and Ebola virus [38] by stimulating the innate immune response. Furthermore, Idarubicin has been shown to be a broad-spectrum enterovirus replication inhibitor, by targeting the internal ribosomal entry site (IRES) [36]. Our study adds another target for Idarubicin: the FSE of HCoV-OC43.

Many of the anthracyclines tested share a similar structure (Figure S5). They all contain an identical tetracycline backbone, with only a few variations of the functional groups. For instance, the only difference between Idarubicin and Valrubicin is a trifluoroacetaldehyde, making the latter inactive. Idarubicin could thus be a suitable scaffold for future new antivirals targeting HCoV-OC43. Also, the only difference between Idarubicin and Daunorubicin is a lack of the methoxy group, which again shows that small changes in the anthracycline structure can change the efficacy of the drug. Addition of Idarubicin and Daunorubicin, but not Nogalamycin, reduced DMS-mediated methylation of specific bases in the HCoV-OC43 -1 FSE. This different mode of binding could be explained by differences in intercalation: Daunorubicin is a classical intercalator, whereas Nogalamycin is a threading intercalator which has an influence on the elasticity of the compound-bound DNA [72]. Another interesting example is Aclacinomycin A, its structure is slightly different from the other anthracyclines (Figure S5), these changes render it ineffective in affecting -1 PRF in this study but also give it the ability to affect topoisomerase I [73] and prevent topoisomerase II induced DNA damage [74]. All these differences in the structures already have big implications on the mode of action, allowing changes that can be favourable to develop drugs showing improved RNA binding specificity and reduced toxicity.

Seasonal coronavirus are common causes of upper respiratory tract infections, and are frequently detected together with other pathogens, with co-infections being associated with increased severity [75]. Re-infections occur frequently, even within six months, and with homologous types such as HCoV-OC43 [76]. HCoV-OC43 infection is associated with more symptomatic infections and remarkably longer duration of illness than the other seasonal CoVs and corresponds to higher viral load. Elderly people with chronic obstructive pulmonary disease (COPD) are at risk for severe, acute disease, exacerbation of COPD and hospitalization when infected by HCoV-OC43 [77]. Moreover, seasonal CoVs cause severe disease in immunocompromised, including cancer patients [78]. Our findings and those of others in the field, that anti-cancer drugs do seem to have an antiviral side to them could be of relevance for these patients. Currently there are studies undertaken to reduce the toxicity of anthracyclines by for example use of combinatory therapies [79]. Further studies in that regard could help to alleviate the potential indirect effect on translation by the anthracyclines to make them a safer antiviral drug.

## Acknowledgments

We thank Baptiste Bogard and Devendra Kumar Maurya for help with the DNA Damage assay and immunofluorescence microscopy. D.S. was funded by the Kempe foundation: JCK22-0048. J.J. was funded by the Swedish Research Council grant #2023-02679, Umeå University, the Stiftelsen Olle Engkvist Byggmästare, Vinnova grant 2019-05491, Chemical Biology Consortium, Sweden and the Erling-Persson Foundation. N.A. was funded by The Swedish Research Council, Dnr: 2023-01831 and 2019-01472; The Swedish Cancer Society, Dnr 22 2005 Pj and 25 4731 Pj; Heart- and Lung Foundation, Dnr 20240660 and 2023-03-08, The Kempe Foundations, Dnr 324325, and SciLifeLab Dnr PLPTDP25:G:014.

## Author contributions: CRediT

DS: Conceptualization, Data curation, Formal analysis, Investigation, Methodology, Validation, Visualization, Writing – original draft, Writing – review and editing.

KI: Conceptualization, Data curation, Formal analysis, Investigation, Methodology, Validation, Visualization, Writing – review and editing

KV: Methodology, Investigation, Writing – review and editing

DI: Formal analysis, Visualization, Writing – review and editing

LP: Methodology, Writing – review and editing

LL: Methodology, Investigation

NA: Conceptualization, Resources, Writing - Review & Editing, Supervision, Project administration, Funding acquisition

JJ: Conceptualization, Resources, Writing - Review & Editing, Supervision, Project administration, Funding acquisition

