## Supplemental Tables 1 and 2, Supplemental figures 1-7 for "Double-edged swords: Anthracyclines inhibit -1 programmed ribosomal frameshifting and restrict HCoV-OC43 infection but show cytotoxicity"

**Short title:** anti-cancer drugs show antiviral activity

**Keywords:** HCoV-OC43, Frameshifting, Anthracyclines, Drug-repurposing, RNA-structure

†These authors have contributed equally to this work and share first authorship

### Table S1: Plasmid list

| **Plasmid** | **Relevant characteristics** | **Reference** |
|---|---|---|
| pEX-A258 | Cloning vector; $Ap^r$ | Eurofins Genomics, Germany |
| pEX-HCoV-OC43 | Synthesized HCoV-OC43 Frameshifting element context DNA vector containing a T7 promoter, leader region and Kozak sequence upstream of an N-terminal 3x FLAG-tag followed by a 290 nt linker. The final 115 bp of the C-terminal end of Nsp10, the coding sequence of Nsp11-Nsp12 with the frameshifting site and the pseudoknot structure and 513 bp of Nsp12 after the frameshift site. A SmaI site was added to the end for runoff transcription. | Eurofins Genomics, Germany<br><br>This study, in a similar fashion as [1] |
| pEX-HCoV-OC43-IFC | pEX-HCoV-OC43 with the frameshifting site disrupted by site-directed mutagenesis PCR with IFC primers bringing Nsp12 in frame. | This study, in a similar fashion as [1] |
| pEX-HCoV-OC43-Stop | pEX-HCoV-OC43 with the frameshifting site disrupted by site-directed mutagenesis PCR with Stop primers, generating a stop codon at the slippery site. | This study, in a similar fashion as [1] |
| pEX-SARS-CoV-2 | Synthesized SARS-CoV-2 Frameshifting element context DNA vector containing a T7 promoter, leader region and Kozak sequence upstream of an N-terminal 3x FLAG-tag followed by a 290 nt linker. The final 115 bp of the C-terminal end of Nsp10, the coding sequence of Nsp11-Nsp12 with the attenuator loop, the frameshifting site and the pseudoknot structure and 531 bp of Nsp12 after the frameshift site. A SmaI site was added to the end for runoff transcription. | Eurofins Genomics, Germany<br><br>[1] |
| pEX-SARS-CoV-2-IFC | pEX-SARS-CoV-2 with the frameshifting site disrupted by site-directed mutagenesis PCR with IFC primers bringing Nsp12 in frame. | [1] |
| pEX-SARS-CoV-2-Stop | pEX-SARS-CoV-2 with the frameshifting site disrupted by site-directed mutagenesis PCR with Stop primers, generating a stop codon at the slippery site. | [1] |
| pSGDlucV3.0 | T7 and SV40 promoter containing dual luciferase (Firefly and Renilla) reporter vector with FMDV StopGo sequences flanking MCS; $Ap^r$ | Addgene Plasmid #119760 [2] |
| pSGDluc-HCoV-OC43 | Dual luciferase reporter containing the final 115 bp of the C-terminal end of Nsp10, the coding sequence of Nsp11-Nsp12 with the frameshifting site, the pseudoknot structure and 100 bp of Nsp12 after the frameshift site. | This study, in a similar fashion as [1] |

| | | |
|---|---|---|
| pSGDluc-HCoV-OC43-IFC | pSGDlucV3.0-HCoV-OC43 with the frameshifting site disrupted by site-directed mutagenesis PCR with IFC primers bringing Nsp12 in frame. | This study, in a similar fashion as [1] |
| pSGDluc-HCoV-OC43-Stop | pSGDlucV3.0-HCoV-OC43 with the frameshifting site disrupted by site-directed mutagenesis PCR with stop primers, generating a stop codon at the slippery site. | This study, in a similar fashion as [1] |
| pSGDluc-SARS-CoV-2 | Dual luciferase reporter containing the final 115 bp of the C-terminal end of Nsp10, the coding sequence of Nsp11-Nsp12 with the frameshifting site, the pseudoknot structure and 91 bp of Nsp12 after the frameshift site. | [1] |
| pSGDluc-SARS-CoV-2-IFC | pSGDlucV3.0-SARS-CoV-2 with the frameshifting site disrupted by site-directed mutagenesis PCR with IFC primers bringing Nsp12 in frame. | [1] |
| pSGDluc-SARS-CoV-2-Stop | pSGDlucV3.0-SARS-CoV-2 with the frameshifting site disrupted by site-directed mutagenesis PCR with stop primers, generating a stop codon at the slippery site. | [1] |

#### Table S2: Oligonucleotide list

| **Name** | **Purpose** | **Plasmid** | **Sequence (5‘->3‘)** |
|---|---|---|---|
| HCoV-IFC-fw | mutagenesis forward primer to disrupt the slippery site of HCoV-OC43 and bring the downstream sequence in frame | pEX-HCoV-OC43-IFC pSGDluc-HCoV-OC43-IFC | ATCAAAAGATACTAATTTCTTCAAGCGGGTTCGGGGTACGAGT |
| HCoV-IFC-rev | mutagenesis reverse primer to disrupt the slippery site of HCoV-OC43 and bring the downstream sequence in frame | pEX-HCoV-OC43-IFC pSGDluc-HCoV-OC43-IFC | ACTCGTACCCCGAACCCGCTTGAAGAAATTAGTATCTTTTGAT |
| HCoV-stop-fw | mutagenesis forward primer to introduce a stop codon into the slippery site of HCoV-OC43 | pEX-HCoV-OC43-stop pSGDluc-HCoV-OC43-stop | AAAGATACTAATTTTTTATAAGGGTTCGGGGTACGAGTG |
| HCoV-stop-rev | mutagenesis reverse primer to introduce a stop codon into the slippery site of HCoV-OC43 | pEX-HCoV-OC43-stop pSGDluc-HCoV-OC43-stop | CACTCGTACCCCGAACCCTTATAAAAAATTAGTATCTTT |

| | | | |
|---|---|---|---|
| HCoV-OC43-fw-XhoI | forward primer to amplify the 3'-end of Nsp10, the frameshift element and 100 bp of the coding region of Nsp12 | pSGDluc-HCoV-OC43 | ATAACTCGAGACCACGCGGCAAGTTTGTACAAG |
| HCoV-OC43-rev-BamHI | reverse primer to amplify the 3'-end of Nsp10, the frameshift element and 100 bp of the coding region of Nsp12 | pSGDluc-HCoV-OC43 | ATAAGGATCCACTAGCATTGTAAATATCAAATGCCC |
| SARS-CoV-2-IFC-fw | mutagenesis forward primer to disrupt the slippery site of SARS-CoV-2 and bring the downstream sequence in frame | pEX-SARS-COV-2-IFC pSGDluc-SARS-CoV-2-IFC | AGCTGATGCACAATCGTTCTTCAAGCGGGTTTGCGGTGTAAGT |
| SARS-CoV-2-IFC-rev | mutagenesis reverse primer to disrupt the slippery site of SARS-CoV-2 and bring the downstream sequence in frame | pEX-SARS-COV-2-IFC pSGDluc-SARS-CoV-2-IFC | ACTTACACCGCAAACCCGCTTGAAGAACGATTGTGCATCAGCT |
| SARS-CoV-2-stop-fw | mutagenesis forward primer to introduce a stop codon into the slippery site of SARS-CoV-2 | pEX-SARS-CoV-stop pSGDluc-SARS-CoV-2-stop | GATGCACAATCGTTTTTATAAGGGTTTGCGGTGTAAGTG |
| SARS-CoV-2-stop-rev | mutagenesis reverse primer to introduce a stop codon into the slippery site of SARS-CoV-2 | pEX-SARS-CoV-stop pSGDluc-SARS-CoV-2-stop | CACTTACACCGCAAACCCTTATAAAAACGATTGTGCATC |
| SARS-CoV-2-fw-XhoI | forward primer to amplify the 3'-end of Nsp10, the frameshift element and 100 bp of the coding region of Nsp12 | pSGDluc-SARS-CoV-2 | ATAACTCGAGACCAACTTGTGCTAATGACCCTGTG |
| SARS-CoV-2-rev-BamHI | reverse primer to amplify the 3'-end of Nsp10, the frameshift element and 100 bp of the coding region of Nsp12 | pSGDluc-SARS-CoV-2 | ATAAGGATCCATTGTAGATGTCAAAAGCCC |
| T7_fw | forward primer to amplify the DNA template for *in vitro* transcription of various RNAs | - | TAATACGACTCACTATAGG |

| | | | |
|---|---|---|---|
| pSGDLuc_IV_rev | reverse primer to amplify the DNA template for *in vitro* transcription of the whole dual-luciferase reporter | pSGDluc-HCoV-OC43 pSGDluc-SARS-CoV-2 | TTACACGGCGATCTTGCC |
| T7-slippery-fw | forward primer to amplify the the DNA template for *in vitro* transcription of the HCoV-OC43 frameshift element, beginning at the slippery site | - | ACATAATACGACTCACTATAGGGCTAATTTTTAAACGGGTTCG |
| Pseudoknot-iv-rev | reverse primer to amplify the DNA template for *in vitro* transcription of the HCoV-OC43 frameshift element, until the end of the pseudoknot structure | - | CAAATGCCCTTAATTGTACATCAG |
| T7-fw-fullFSE | forward primer to amplify the the DNA template for *in vitro* transcription of the HCoV-OC43 frameshift element, beginning at 3-end of Nsp10 | - | ACATAATACGACTCACTATAGGGCGCGGCAAGTTTGTACAAGTG |
| fullFSE-iv-rev | reverse primer to amplify the DNA template for *in vitro* transcription of the HCoV-OC43 frameshift element, including the 100 bp of Nsp12 | - | CACTAGCATTGTAAATATCAAATGCC |
| 5′RA (Ribooligonucleotid) | 5’ adapter for Illumina TrueSeq adapter ligation | - | UCCCUACACGACGCUCUUCCGAUCU |
| 3′DA | 3’ adapter for Illumina TrueSeq adapter ligation | - | 5phos/AGATCGGAAGAGCACACGTCTGAACTCCAG/3ddC |
| LibrAMP-F | forward primer to amplify Illumina TrueSeq adapter ligated libraries for sequencing | - | AATGATACGGCGACCACCGAGATCTACACTCTTTCCCTACACGACGCTCTTC |
| LibAmp_RPI1_R | reverse primer to amplify Illumina TrueSeq adapter ligated libraries and add an index for sequencing | - | CAAGCAGAAGACGGCATACGAGATCGTGATGTGACTGGAGTTCAGACGTGTGCT |
| LibAmp_RPI2_R | reverse primer to amplify Illumina TrueSeq adapter | - | CAAGCAGAAGACGGCATACGAGATACATCGGTGACTGGAGTTCAGACGTGTGCT |

| | | | |
|---|---|---|---|
| | ligated libraries and add an index for sequencing | | |
| LibAmp_RPI3_R | reverse primer to amplify Illumina TrueSeq adapter ligated libraries and add an index for sequencing | - | CAAGCAGAAGACGGCATACGAGATTTAGGCGTGACTGGAGTTCAGACGTGTGCT |
| LibAmp_RPI4_R | reverse primer to amplify Illumina TrueSeq adapter ligated libraries and add an index for sequencing | - | CAAGCAGAAGACGGCATACGAGATTGGTCAGTGACTGGAGTTCAGACGTGTGCT |
| LibAmp_RPI5_R | reverse primer to amplify Illumina TrueSeq adapter ligated libraries and add an index for sequencing | - | CAAGCAGAAGACGGCATACGAGATCACTGTGTGACTGGAGTTCAGACGTGTGCT |
| LibAmp_RPI6_R | reverse primer to amplify Illumina TrueSeq adapter ligated libraries and add an index for sequencing | - | CAAGCAGAAGACGGCATACGAGATATTGGCGTGACTGGAGTTCAGACGTGTGCT |
| LibAmp_RPI7_R | reverse primer to amplify Illumina TrueSeq adapter ligated libraries and add an index for sequencing | - | CAAGCAGAAGACGGCATACGAGATGATCTGGTGACTGGAGTTCAGACGTGTGCT |
| LibAmp_RPI8_R | reverse primer to amplify Illumina TrueSeq adapter ligated libraries and add an index for sequencing | - | CAAGCAGAAGACGGCATACGAGATTCAAGTGTGACTGGAGTTCAGACGTGTGCT |
| LibAmp_RPI9_R | reverse primer to amplify Illumina TrueSeq adapter ligated libraries and add an index for sequencing | - | CAAGCAGAAGACGGCATACGAGATCTGATCGTGACTGGAGTTCAGACGTGTGCT |
| LibAmp_RPI10_R | reverse primer to amplify Illumina TrueSeq adapter ligated libraries and add an index for sequencing | - | CAAGCAGAAGACGGCATACGAGATAAGCTAGTGACTGGAGTTCAGACGTGTGCT |
| Enrich_F | forward primer to amplify Illumina TrueSeq adapter ligated libraries for sequencing | - | AATGATACGGCGACCACCGAGATC |
| Enrich_R | reverse primer to amplify Illumina TrueSeq adapter ligated libraries for sequencing | - | CAAGCAGAAGACGGCATACGAGAT |

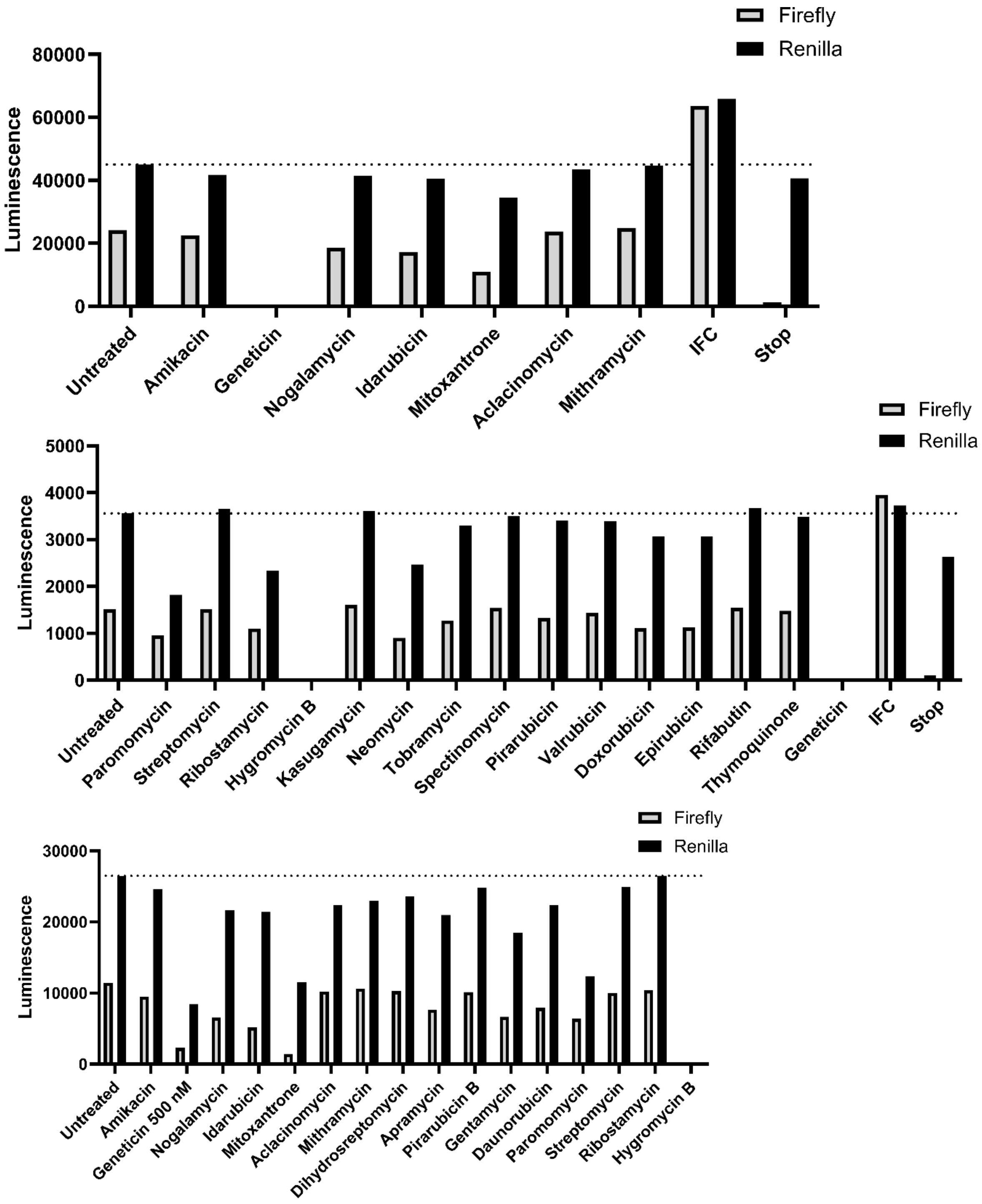


**Figure S1. General translation inhibiting effect of several compounds.** Raw luminescence data for some of the tested compounds as depicted by representative experiments. The concentration of the different compounds is 10 µM unless otherwise stated. Some of the tested compounds (e.g. Geneticin) exhibited severe general translation inhibition in the *in vitro* translation system.

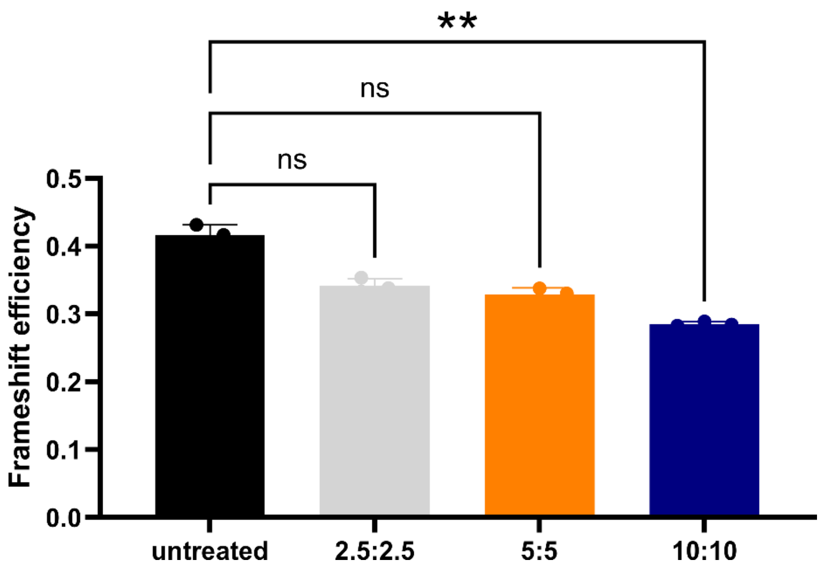


**Figure S2. Nogalamycin and Idarubicin show no synergistic effect on frameshift inhibition.** *In vitro* translation of the Dual luciferase-HCoV-OC43-FSE reporter by rabbit reticulocyte lysate. Nogalamycin and Idarubicin were added in an equal ratio at the indicated concentrations to the translation mixture. Experiments were carried out three times. Mean and corresponding standard deviation are shown. The *p*-values are indicated as $^{**}p<0.01$; ns, not significant.

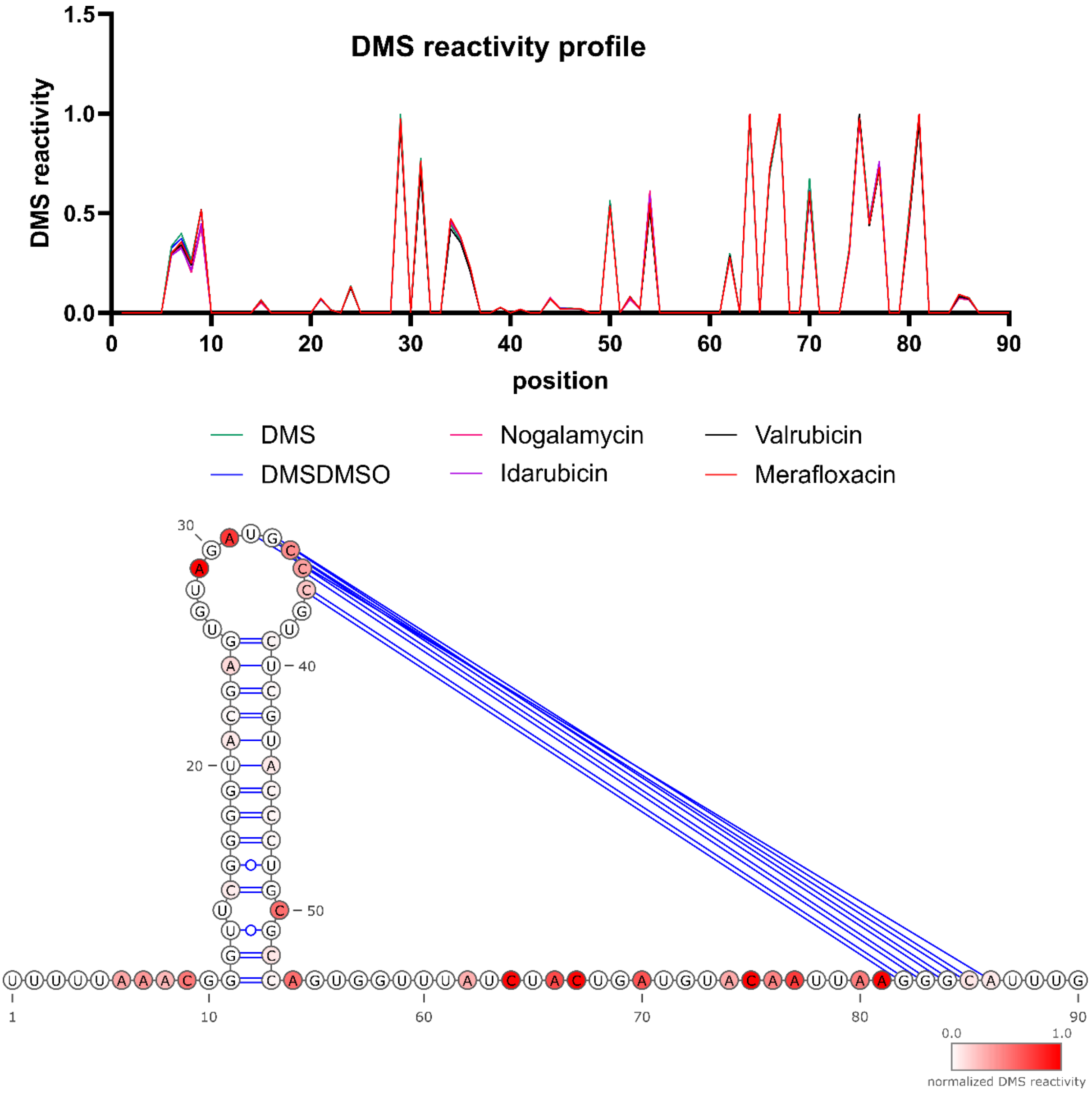


**Figure S3. DMS reactivity is not changed when compounds are added post-folding. Upper panel:** Normalized DMS reactivities in absence or presence of compound. **Lower panel:** Resulting secondary structure of FSE deducted from the DMS-reactivity.

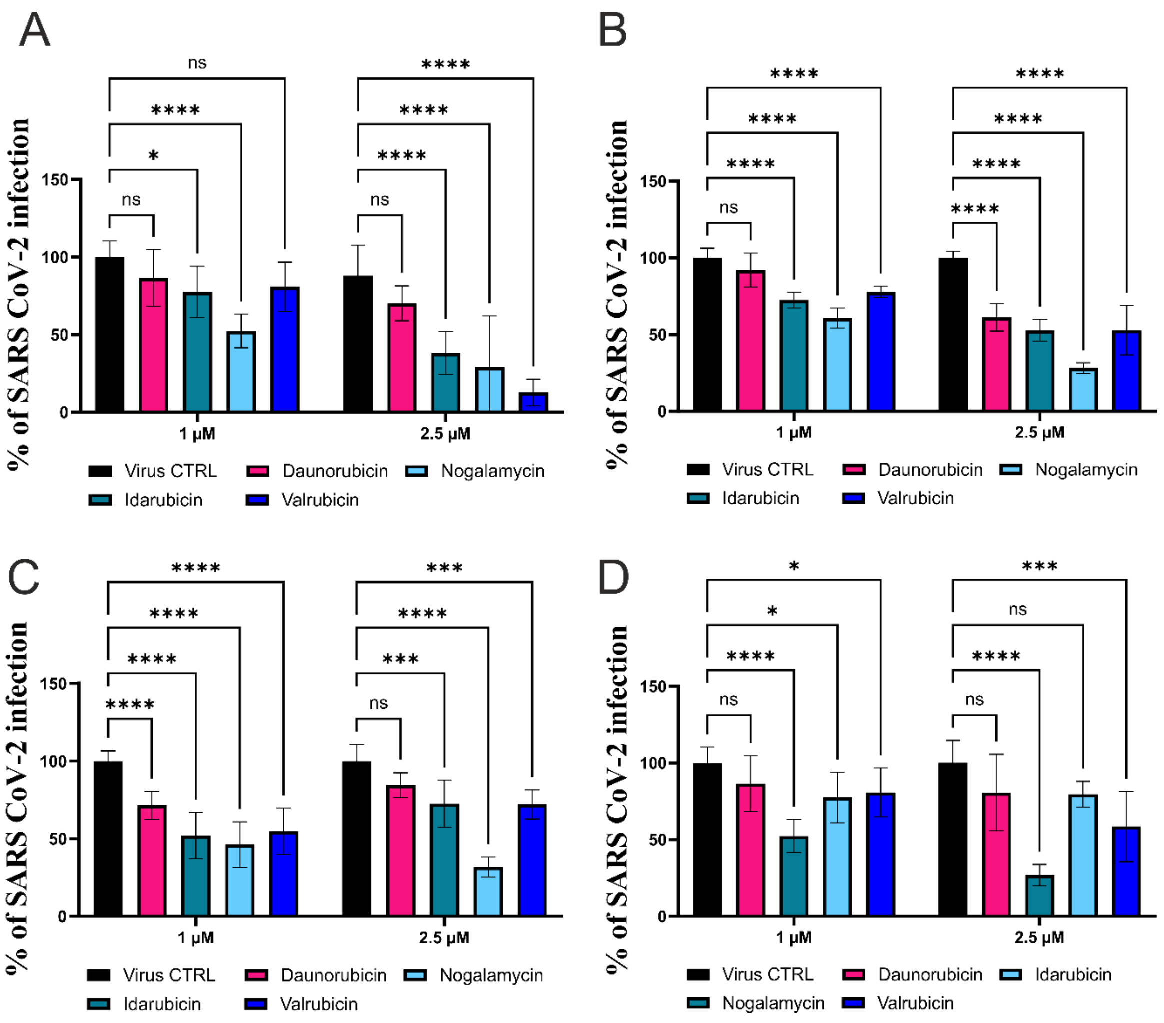


**Figure S4. Anthracyclines reduce SARS-CoV-2 infection.** Inhibition of SARS-CoV-2 strain **(A)** Wuhan **(B)** Omicron and **(C)** XBB.1.9.2.5.1.3 infection by anthracyclines. A549-ACE2 cells were infected with SARS-CoV-2 (MOI =0.1) for 16h in presence of 1 and 2.5 µM of the indicated compound. The viral titer was determined by immunofluorescence with a nucleocapsid antibody. Results were generated with three (1 µM) or two (2.5 µM) independent experiments, each with three replicates. Mean and corresponding standard deviation of all replicates are shown. *p*-values are indicated as **p*<0.05; ***p*<0.01; ****p*<0.001; *****p*<0.0001; ns, not significant.

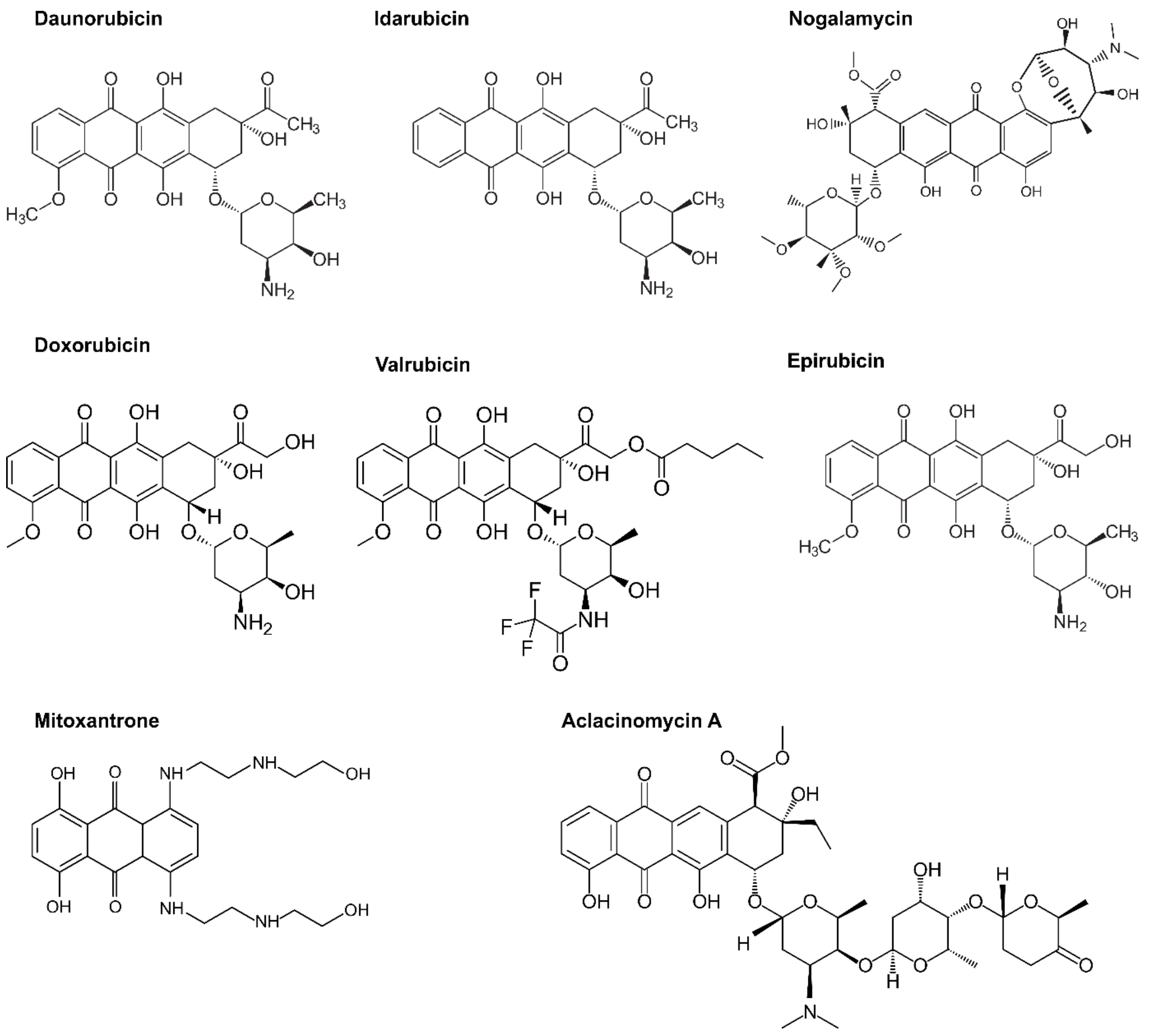


**Figure S5. Chemical structures of tested anthracyclines.**

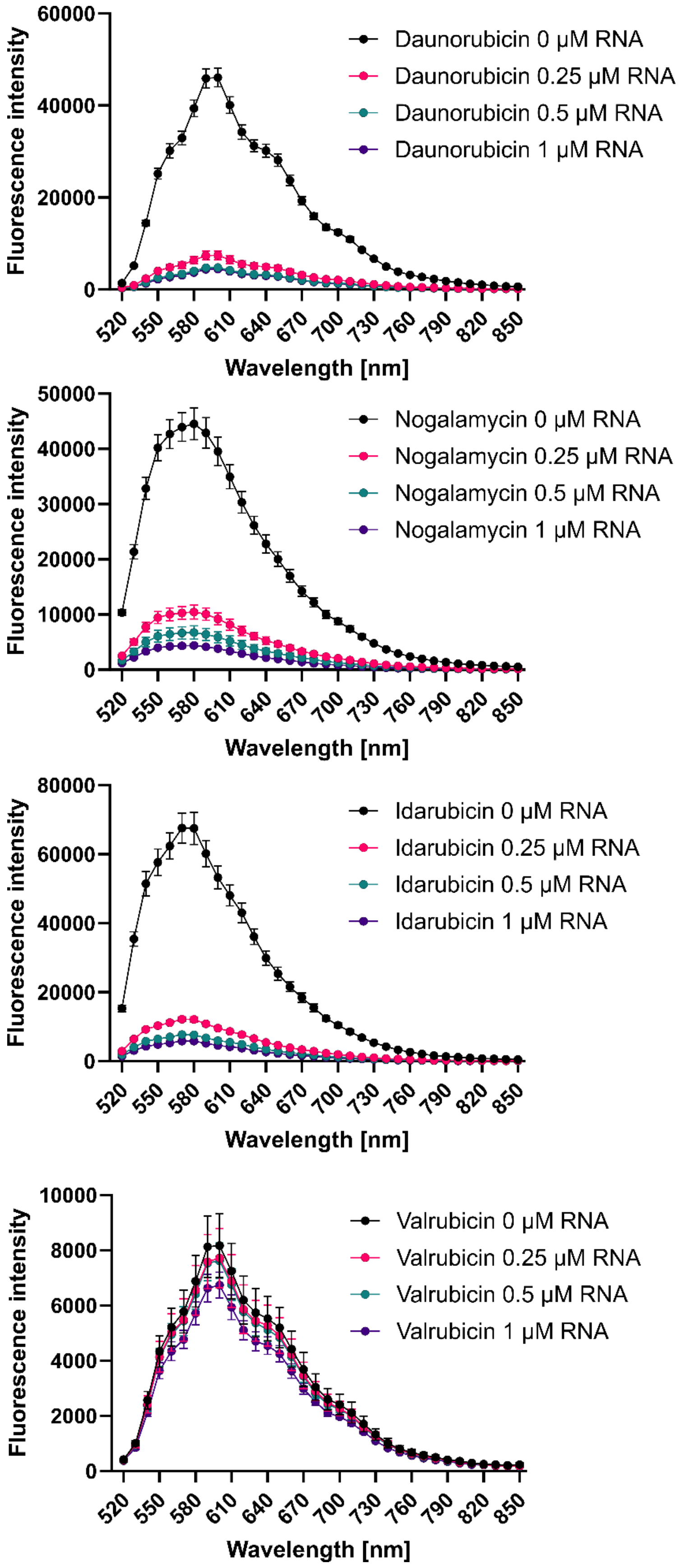


**Figure S6. Emission spectra recorded after excitation at 482 nm for the different compounds in absence and presence of** SARS-CoV-2 FSE RNA.

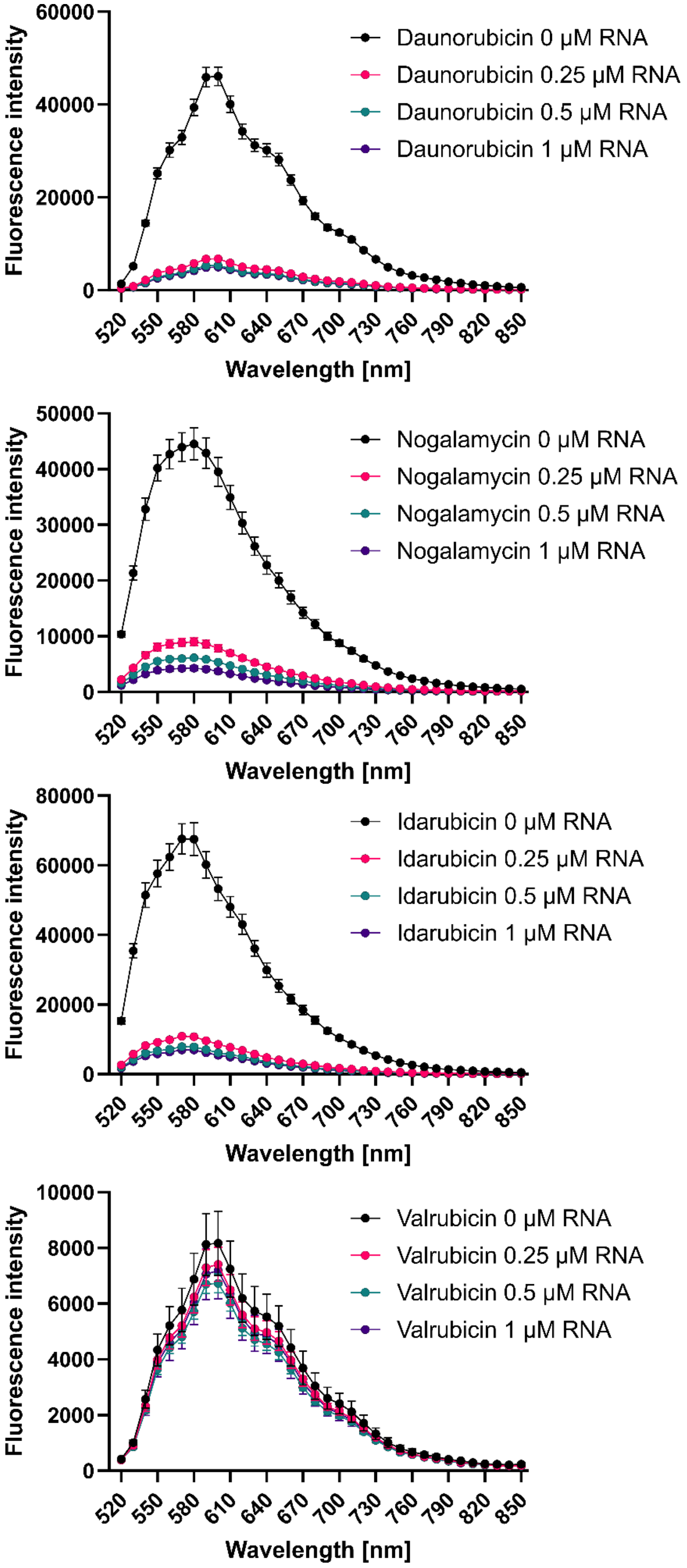


**Figure S7. Emission spectra recorded after excitation at 482 nm for the different compounds in absence and presence of** bacterial *prfA*-RNAT RNA.
